# A ribosomal protein variant that confers macrolide resistance differentially regulates acid resistance, catabolism, and biofilm formation related genes in *Escherichia coli*

**DOI:** 10.1101/2021.04.10.439278

**Authors:** Elizabeth A. Franklin, Sarah B. Worthan, Chi Pham, Mee-Ngan F. Yap, Luis R. Cruz-Vera

## Abstract

Mutational changes in bacterial ribosomes that confer antibiotic resistance decrease cell fitness. Determining the genetic factors that interconnect antibiotic resistance and cell fitness is critical in the fight against bacterial infections. Here, we describe gene expression and phenotypic changes presented in *Escherichia coli* cells carrying an uL22(K90D) mutant ribosomal protein, which showed growth defects and resistance to macrolide antibiotics. Ribosome profiling analyses revealed reduced expression of operons involved in catabolism, electron transportation, indole production, and lysine-decarboxylase acid resistance. In general, ribosome occupancy was increased at rare codons while translation initiation of proximal genes in several of the affected operons was substantially reduced. Decline of the activity of these genes was accompanied by increased expression of macrolide multidrug efflux pumps, the glutamate-decarboxylase regulon, and the autoinducer-2 metabolic regulon. In concordance with these changes, uL22(K90D) mutant cells grew better in acidic conditions and generated more biofilm in static cultures than their parental strain. Our work provides new insights on how mutations in ribosomal proteins induce the acquisition of macrolide and pH resistance, and increase the ability to generate biofilms.

## INTRODUCTION

All actively translating nascent peptides must traverse the ∼100 Å long ribosomal exit tunnel during their synthesis. The ribosomal exit tunnel is composed of both ribosomal proteins and rRNA residues, which contribute to the architecture and irregular shape of the tunnel, resulting in a cavity that contains bumpy and pitted walls with areas of varying widths [1]. The narrowest region of the tunnel, named the constriction region, is comprised of 23S rRNA nucleotides and the extended loops of the two riboproteins uL4 and uL22 [1]. The constriction region separates the tunnel into two segments, an upper and lower segment. The upper segment, located between the ribosome active site known as the peptidyl transferase center (PTC) and the constriction region, is about 50 Å long [1,2] and can accommodate between 12-16 amino acid residues of a full extended or alpha helical nascent peptide during translation [2]. The upper segment of bacterial ribosomes is less variable than the lower segment [1] and is known to contain binding sites for antibiotics [3], small metabolites, and nascent peptides that are involved in regulating the progress of translation [2]. Passing the constriction region, the lower segment of the tunnel widens, allowing nascent peptides to acquire initial folding structures [4]. Due to its structural location, the constriction region is considered a gateway between the two functional upper and lower segments of the exit tunnel [5].

Changes in the components constituting the upper segment of the tunnel and constriction region affect the translational regulatory function of antibiotics, small molecules, and nascent peptides [2,3,6]. Resistance mutations to sub-inhibitory macrolide antibiotic concentrations often occur in the upper segment of the tunnel [3] and are frequently mapped to the uL4 and uL22 ribosomal proteins that line the tunnel wall (**Figure 1A**) [7,8]. These mutational changes, which could be single or multiple, confer an increased likelihood of obtaining additional mutations in the ribosome or other genes, boosting survival under high antibiotic concentrations but producing variable costs in cellular fitness [9-11]. Mutations in uL4 and uL22 ribosomal proteins also affect the function of regulatory nascent peptides that interact with small molecules in the exit tunnel, controlling protein synthesis and gene expression [2]. An example of such mutational effects is seen during the regulation of the *tnaCAB* (*tna*) operon, which is responsible for the breakdown of L-tryptophan (L-Trp) to indole [12]. The *tna* operon is controlled by a dedicated transcriptional attenuation mechanism that senses free L-Trp [13]. Free L-Trp interacts at the constriction region of ribosomes translating the *tna* leader peptide, TnaC, inducing translation arrest and allowing the transcription of the downstream genes expressing tryptophanase (TnaA) and a L-Trp permease (TnaB) [13]. Mutational changes in the uL22 K90 residue abolish L-Trp-induced translation arrest at the *tnaC* gene and as a consequence the expression of the *tna* operon and production of indole [14]. Well characterized uL4 and uL22 erythromycin resistant mutations have shown different global translational effects as well. A mutation in uL4 that changed a highly conserved residue was associated with a reduction of erythromycin binding to ribosomes and multiple translation defects [15]. In this same study, another mutation conferring erythromycin resistance, which generates a deletion in the extended loop of uL22 (Δ82-84) and widens the constriction region [16] without affecting erythromycin binding to ribosomes [7], was found not to promote decoding errors [15]. However, a recent study determined that this uL22 (Δ82-84) mutational change affects expression of the *tna* operon, functioning of the Sec secretory system, and consequently the expression of several genes related to bacterial virulence and survival [17]. Overall, these observations indicate that changes in the uL4 and uL22 ribosomal proteins that generate resistance to antibiotics affect translation progression, expression of genes, and thus cellular fitness [17]. Furthermore, studies on regulatory nascent peptides have shown that sequence variations among uL22 homologs exhibit species-specific expression of genes [6], suggesting that single mutational changes in uL22 could alter global expression of genes in bacteria.

**Figure 1.**
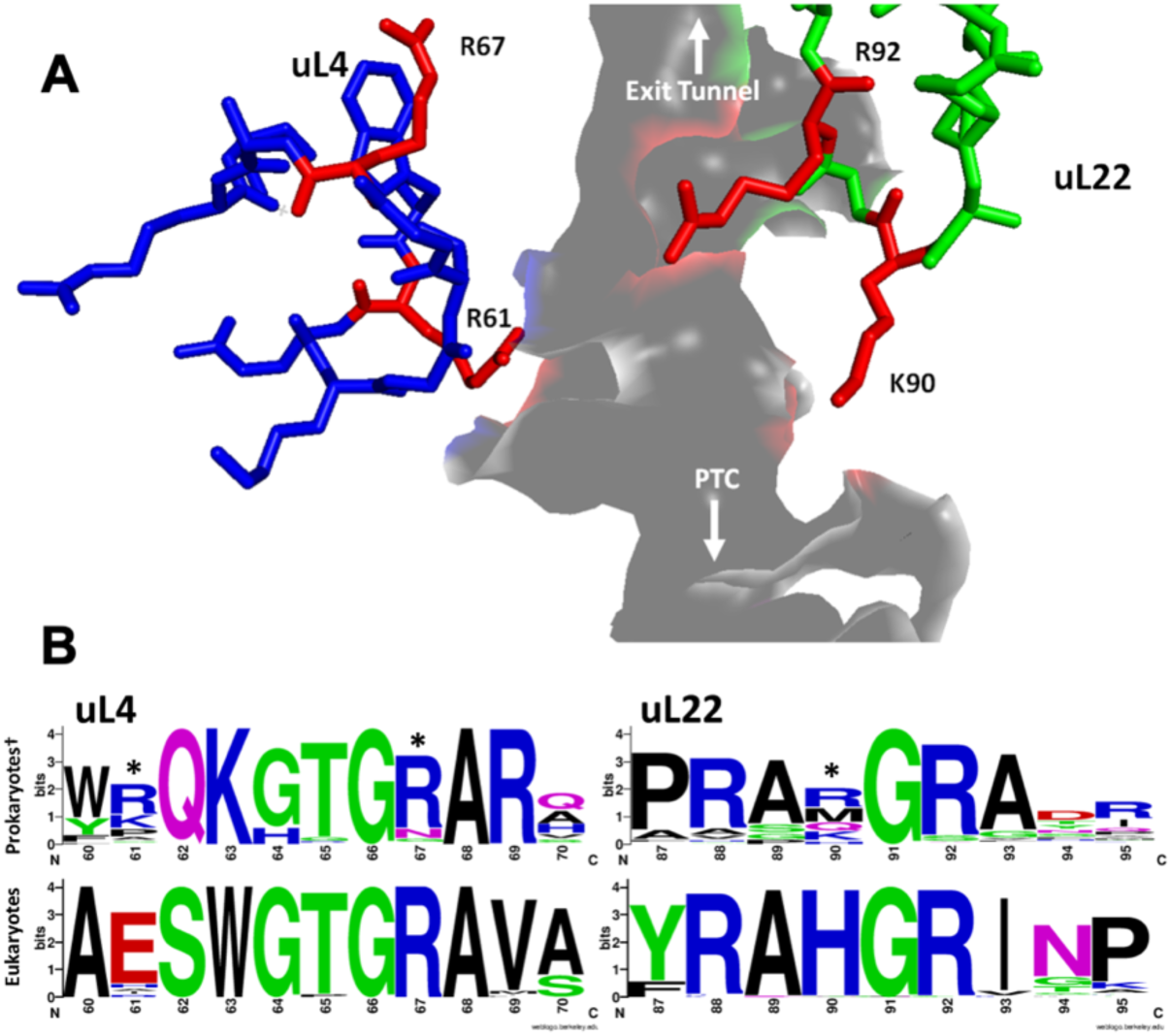
Comparison analysis of uL4 and uL22 amino acid sequences from diverse prokaryotic and eukaryotic organisms. (A) In this figure obtained from the PDB:4UY8 structure, the structural extended loops of the ribosomal proteins uL4 (blue sticks) and uL22 (green sticks) are surrounding at the constriction region the lumen of the ribosomal exit tunnel (gray surface). Residues of both proteins, uL4 R61 and R67, and uL22 K90 and R92, pointing toward the growing peptide chain are shown in red. The orientation of the PTC and the tunnel exit are indicated with arrows. (B) Weblogos results of multiple protein alignment analysis of segments of the extended loops of uL4 and uL22 proteins that constitute the constriction region of the ribosomal exit tunnel. These four Weblogos represent the conservation and frequency of amino acid residues at specific positions *(E. coli* numbering). Prokaryotic sequences are shown on top (231 species), and eukaryotic sequences are shown on bottom (56 species). Residue positions studied in this work are marked with an asterisk. ^†^No archaea sequences were included in this alignment.

In this work, we set out to determine the effects of single residue changes in non-conserved positions of uL4 and uL22 on the global expression of genes. Initially, growth curves were conducted in different conditions to assess the *in vitro* cellular fitness of *E. coli* harboring various mutational changes in uL4 and uL22. We also evaluated the sensitivity of uL22 mutations to various classes of antibiotics. The mutation uL22(K90D) reduced cell growth and antibiotic sensitivity, making it an ideal candidate to determine the effect of this mutation on global gene expression. Ribosome profiling analysis indicated alterations in levels of mRNA transcripts for ∼ 7% of genes and changes in the translational efficiency in 1% of genes expressed. In general, genes related to metabolism of alternative carbon sources, amino acids, and respiration showed a reduction in translational efficiency, as well as in their mRNA levels. Particularly, two genes related to acid resistance response, *cadB* and *adiY*, display the highest reduction in their translational efficiency. On the other hand, translational efficiency of several genes related to ABC transporters was significantly increased. Additionally, there were increases in the mRNA abundance of genes involved in the glutamate-dependent acid resistance response and in the uptake/modification of the autoinducer-2 (AI-2) signaling molecule. These increases correlated with the ability of the cells to resist acidic changes in their growth media and to produce biofilms. In summary, we found general alterations in translation initiation and progression of several genes that pinpoint possible candidate genes whose translation could be affected directly by the ribosomes harboring the uL22(K90D) mutational change and these expressional changes may induce compensatory alterations in the expression of other genes.

## MATERIALS AND METHODS

### Molecular modeling and weblogo creation

The structure of the *E. coli* 50S ribosomal subunit (PDB 4UY8) containing a regulatory nascent peptide was modeled using PyMOL (The PyMOL Molecular Graphics System, Version 2.0 Schrödinger, LLC). Weblogos were generated as previously described [18,19]. Extended loop sequences for uL4 and uL22 homologous proteins were aligned from 231 species across 16 bacterial phyla and for 56 species across 18 eukaryotic phyla. Archaea were excluded from the alignments. Prokaryotic phyla with less than 10 species were not included in these figures and no partial, multispecies, or consensus sequences were used.

### Bacterial strains, plasmids, and growth media

A full list of bacterial strains and plasmids used in this work can be found in **Supplemental Table 1**. Luria Broth (LB) (5 g/L sodium chloride, 10 g/L tryptone, 5 g/L yeast extract), broth and agar (15 g/L), was used as rich media to maintain and test the studied strains. Minimal M9 media (M9) (140 mM disodium phosphate, 20 mM potassium monophosphate, 8 mM sodium chloride and 18 mM ammonium chloride, 2 mM magnesium sulfate, and 0.1 mM calcium chloride) supplemented with 0.2% glycerol and 0.05% casamino acids, unless otherwise indicated, was used to test bacterial growth. Diluted LB media was prepared with sterile deionized water. Acidic and basic media was prepared by the addition of either HCl or KOH to LB media until the desired pH was reached. The media was then sterilized via filtration through a 0.2 μm Nalgene syringe filter (Thermo SCIENTIFIC). Antibiotic stock solutions were prepared by dissolving each antibiotic in the appropriate solvent and stored at -20°C, with dilutions prepared in sterile deionized water. For maintenance and testing, the following concentrations of antibiotics were used: 10 µg/ml tetracycline (Tet) and 50 µg/ml kanamycin (Km).

### Generation of uL4 and uL22 mutant strains

Briefly, *E. coli* K-12 strain SVS1144 was used to generate uL4 and uL22 mutant strains. Initially, site-directed mutagenesis was performed targeting desired codon positions of the *rplD* and *rplV* genes on the pS10 plasmid [20] using the QuikChange Lightning kit (Agilent Technologies) and the primers indicated in **Supplemental Table 2**. Mutated plasmids were Sanger-sequenced (Eurofins) to confirm the desired mutation. Mutant plasmids were transformed into SVS1144. To eliminate expression of the chromosomal *s10* operon, which expresses both uL4 and uL22 wild type proteins, pS10 derived transformed strains were P1vir transduced with viruses obtained from the SM1110 strain [20]. Resulting transductants were growth-tested in LB media plates with Km and Tet antibiotics to confirm the chromosomal insert of the Kan^R^ cassette and retention of the pS10 plasmid. Thereafter, transduced strains were grown for further experiments without Km and Tet antibiotics as suggested previously [20].

### Growth Curves

Cellular growth was measured using a Klett photoelectric colorimeter, Manostat (Cat #76-500-000). This instrument measures the intensity of scattered light, thus allowing for comparison of the density of cells suspended in liquid media. Cultures were grown in a nephelo culture flask using LB, diluted LB, and M9 media with and without casamino acids supplementation. Prior to the start of each experiment, the OD_600_ of overnight cultures was measured and normalized according to cell density. Cell growth was monitored and expressed as Klett units as a function of time.

### Calculation of antibiotics minimal inhibitory concentration (MIC)

MIC experiments by broth microdilution were performed as described previously [21] with some modifications. Cells were grown overnight in LB media, diluted, and added to the appropriate well in a 96-well plate with prepared concentrations of the specified antibiotic. These cells were incubated with shaking in a microplate reader (SpectraMax iD5) for 12 hours at 37°C and the OD_600_ was measured periodically. MIC values correspond to the antibiotic concentrations where the OD_600_ absorbance values detected at any time during the period of incubation were not different from the starting OD_600_ absorbance values.

### Ribosome Profiling

Cells were subcultured from an overnight culture and grown in 100 mL of LB media to an approximate OD_600_ absorbance of 0.6 (corresponding to 120-160 Klett units). 100 µg/mL of chloramphenicol (Cm) was then added and cultures were left to grow for three more minutes. Cells were harvested via centrifugation (4,500 x g for 10 minutes at 4°C) with a Sorvall LYNX 6000 Superspeed Centrifuge. Cell pellets were resuspended in 5 mL 1X Polysome Buffer (20 mM Tris-HCl pH 7.4, 100 mM sodium chloride, 5 mM magnesium chloride) supplemented with 100 µg/mL Cm and centrifuged again (3,220 x g for 10 minutes at 4°C) using an Eppendorf 5804 benchtop centrifuge. The liquid was then decanted and pellets were flash frozen using liquid nitrogen. RNA sample preparations were performed as described previously [22] using the ARTseq™ and Epicentre’s Ribo-Zero™ kits (Illumina). Briefly, samples were lysed using the kit provided polysome buffer and polysomes were digested with micrococcal nuclease. The monosome fraction was obtained by using a size-exclusion chromatography column and RNA was then extracted from the monosome fragment. rRNA was depleted by using the Ribo-Zero kit and PAGE purification of the ribosome protected fragments (RPFs), between 15 – 50 nt, was followed by cDNA synthesis. All RIBO-seq library preparation and sequencing were performed by TB-SEQ, Inc. with an Illumina NovaSeq6000 sequencer. Both raw and trimmed (of adaptor sequences) RNA-seq and RIBO-seq libraries were generated and quality control data was provided for each sample in both library sets.

### Ribosome Profiling Analysis

First, all trimmed fastQ sequences were confirmed to correspond to the appropriate wild type or uL22(K90D) cell samples using the Grep command in Linux. The trimmed sequences were then processed using two different pipelines. The RiboGalaxy and RiboToolkit web tools were used to visualize and quantify the RNA-seq and ribosome profiling sequence data, respectively [23,24]. Each tool processed the same raw data, with the qualitative read count visualization from RiboGalaxy matching the quantitative outputs of RiboToolkit. Adapter trimmed fastq sequences were first uploaded to RiboGalaxy and the read quality was double-checked using FastQC [25]. All sequence data displayed high quality reads up to 40 bp (data not shown). Once quality was confirmed, rRNA was removed from the samples using Bowtie [26] and then the remaining reads were aligned to the provided reference genome. The default settings were used as well as the *E. coli* str. K-12 substr. MG1655: eschColi_K12 reference index and genome. The aligned file output was converted from SAM to BAM format and then a ribosome profile (GWIPS-viz Mapping tool) was generated using an offset from the 3’-end with default settings corresponding to an A-site (of 12) offset and the eschColi_K12 reference genome (GenBank # GCA_000005845). The resulting bigwig files were visualized through GWIPS-viz [27]. Generated BAM files of each sample replicate were then merged (summed) to combine the data into a single track using RiboGalaxy and then visualized using the Integrative Genomics Viewer (IGV) [28].

RiboToolkit is an online resource for analyzing RNA- and Ribo-seq data. Manipulating the data for input was done on an Ubuntu 20.04.1 LTS server accessed remotely using PuTTY with all necessary packages, libraries, and dependencies downloaded directly to the server. Genome assembly ASM584v2 was downloaded from https://bacteria.ensembl.org and used as a reference genome for alignment [29]. Inputs for RiboToolkit were produced by following the suggested preparations (sequences were already trimmed and cleaned upon receipt). The utility to create the collapsed FASTA files was downloaded and implemented through perl to generate the inputs for the single case ribo-seq analysis. The featureCounts utility from the subread package was used to create a gene count matrix of the RNA counts of all samples to generate the input for the group case study [30,31]. The BAM files for input into featureCounts were generated using Bowtie (1.2.3) default settings for read count alignments and samtools (1.10) for SAM to BAM format conversion [32]. *E. coli* K-12 was used as the selected species for all analysis/tools in RiboToolkit. Single case analysis of each ribo-seq sample was performed using the default settings except for RPF length, which was broadened to 20 (shortest) and 40 (longest). Next, group analysis was performed using all the previous single case results combined with the RNA-seq data to yield information about differential translation between groups using default settings. The averages of the read counts for both mRNA and RPFs for each sample group (wild type or K90D) were determined. Fold change in mRNA abundance and ribosome occupancy were calculated as the ratio mRNA of uL22(K90D)/ mRNA of wild type, and RPFs of uL22(K90D)/ RPFs of wild type average read counts for each gene, respectively. These ratio values were also used to determine the adjusted p-values (corrects for false discovery rate). Translation efficiency was calculated as the ratio RPFs/ mRNA read counts per each gene. Fold change in translation efficiency was calculated as the ratio translation efficiency of uL2(K90D)/ translation efficiency of wild type values [24]. The additional codonstat tool was also used to examine specific codon frequency in genes indicated as having altered translation efficiency based off of A-, P-, and E-site codon occupancy indicated in the group analysis, background setting was changed to random-select and codons to all codon.

### Biofilm Assays

Biofilm formation assays were performed as previously described [33]. Starter cultures were grown overnight at 37°C for 16-18 hours and then diluted 1:1000 in LB media. One mL of diluted overnight culture was added to 14 mL polypropylene round-bottom tubes (FALCON) and incubated at 37°C in a static upright position for 24 hours. Following incubation, planktonic cells were removed from the tubes by a series of three washes with deionized water. After tubes were air-dried, 1.5 mL of 0.1% Gram Crystal Violet (Remel Inc) was added and used to stain adherent cells for 15 minutes. The crystal violet solution was removed and the tubes were washed once more with deionized water three times and left to air-dry. Once dry, 1.5 mL of an ethanol:acetone (80:20) solution was added to tubes as a solubilizing agent and left at room temperature for 15 minutes. The absorbance of the solution was measured in duplicate for each tube at a wavelength of 570 nm (BIO-TEK PowerWaveHT).

## RESULTS

### Variability of the uL4 and uL22 amino acid residues that constitute the constriction region of the ribosomal exit tunnel

Previous structural and computational studies of ribosomes indicate that specific residues of uL4 and uL22 extension loops are in the path of growing nascent chains (**Figure 1A**) [2,34]. It has been suggested that interactions between these ribosomal proteins and nascent peptides possibly affect the rate of translation elongation/termination and support initial folding of nascent peptides [2,35], thus potentially having the ability to modulate gene expression and protein production.

Given that previous analysis has suggested the variability of uL4 and uL22 protein residues among bacteria is determined by the need for species-specific protein synthesis regulation [6] and because we were interested in understanding the role uL4 and uL22 play in global gene expression, we initially decided to determine the chemical conservation of the residues constituting the extended loops of these ribosomal proteins by comparing their orthologs among prokaryotic and eukaryotic organisms. For bacterial species, we compared 231 sequences spanning sixteen phlya using a weblogo generator that creates sequence logos based on multisequence alignments and displays the conservational frequency of an amino acid at a given position (**Figure 1B**). We noted that in prokaryotes while many positions were highly conserved, likely due to important structural interactions or functions, others were more variable. These extended loops sequences show an enrichment of basic amino acids and an absence of acidic residues (**Figure 1B**), as previously observed [1]. In particular, many of the conserved basic amino acids in these loops point inward toward rRNA. The majority of the residues pointing outward into the constriction region of the tunnel, toward the growing nascent peptide, were variable. Variability was observed in K90 of uL22 and R61 and R67 of uL4, with the majority of orthologous proteins containing basic amino acids. Only the R92 residue in uL22, while basic as well, was invariant (**Figure 1B**). Similar observations were noted for eukaryotic uL4 and uL22 residues. Although in eukaryotes the overall conservation of residues is more prominent compared to prokaryotes, there was an analogous enrichment of basic amino acids, especially in residues which face the tunnel, with the exception of a glutamic acid residue at the 61st position in uL4 (**Figure 1B**). Overall, our in-silico analyses indicated that, in prokaryotes, uL4 and uL22 residues pointing towards the exit tunnel are frequently variable. However, despite their variability, there is a general avoidance of negatively-charged amino acids in their extended loops.

### Effects of replacing non-conserved amino acid residues of the uL4 and uL22 extended loops on cell growth

To investigate the importance of the observed avoidance of acidic amino acids at non-conserved residue positions of ribosomal proteins that constitute the constriction region on bacterial cell fitness, the uL4 R61, R67 and uL22 K90 residues were mutated to an aspartic acid. Growth curves were performed in rich media (LB) or minimal media (M9 with any supplements indicated) to differentiate between any possible nutritionally dependent effects (**Materials and Methods**). We did not observe any significant changes in the growth behavior of cells with the uL4(R61D) or uL4(R67D) mutations with respect to the wild type (**Supplemental Figure 1**). However, we did observe that uL22(K90D) bacterial cultures consistently display a slower growth phenotype than their parental bacterial cultures (**Figure 2A**). uL22(K90D) bacterial cultures also exited exponential growth, preparing for stationary phase entry, at a lower cellular density than uL22 wild type cultures (**Figure 2A**). The slow growth phenotype is first evident at the onset of exponential phase (**Figure 2A, arrows**) and becomes most apparent at the transition between late log and stationary phase. Furthermore, growth in diluted LB media makes the growth differences between uL22(K90D) and wild type strains more apparent (**Figure 2B; highlighted values**). Interestingly, we did not observe significant differences in growth between the uL22(K90D) and the wild type strains when grown in M9 media (**Figure 2B; highlighted values**), which indicates the growth differences observed in LB rich media are dependent on its chemical composition. Thus, the consistent departure from exponential growth at a lower cell density in rich media by cells containing the uL22(K90D) mutant protein indicates a phenotypic change warranting further investigation.

**Figure 2.**
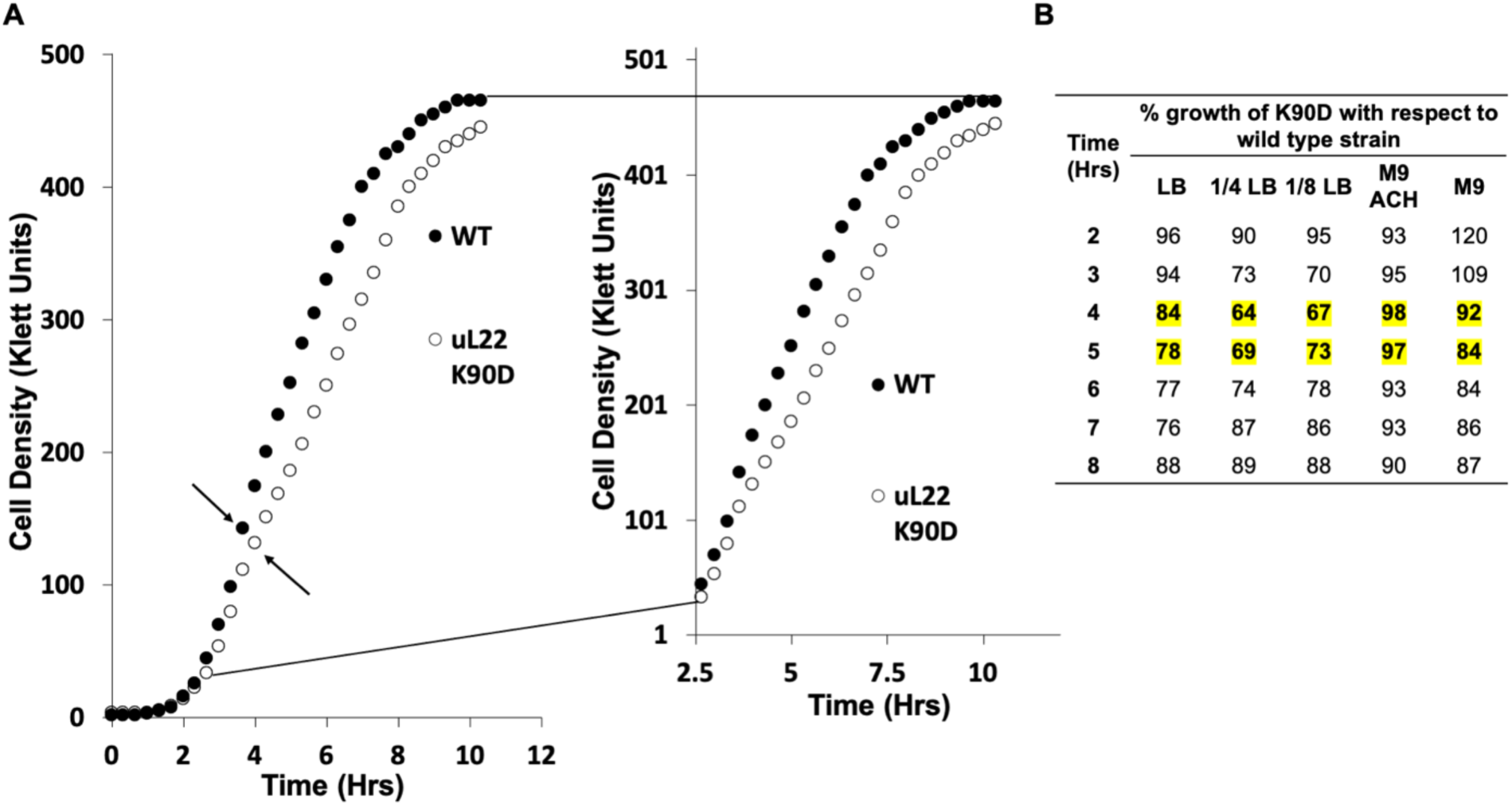
Cell growth of wild type and uL22(K90D) mutant strains in varying media. (A) Plot representing bacterial cultures growth curves of wild type and uL22(K90D) strains. Right panel corresponds to a zoomed view of the resulting curve to highlight the growth differences in the late exponential and entry of the stationary phase. Arrows indicate regions of the curve where samples were taken for ribosome profiling and RNA-seq analyses. This figure is representative of twenty independent experiments. (B) Percentage (%) growth was obtained by dividing the Klett unit density values obtained for cultures of the uL22(K90D) strain over the Klett unit values for cultures of the wild type strain during the indicated times. Highlighted values show the biggest differences in growth among media. These values are representative of at least two independent experiments.

### Effects of the uL4 and uL22 aspartic mutational replacements on antibiotic resistance

Mutations in uL4 and uL22 are known to be associated with increased antibiotic resistance in *E. coli* laboratory isolates [7,8] as well as in field isolates of several species [36–40] and cells containing mutations that confer antibiotic resistance are known to frequently have an associated fitness cost [9-11]. Notably, the tylosin (a macrolide) resistant mutations at position 90 (glutamine to lysine or histidine) in *Mycoplasma agalactiae*, which is relevant to the dairy industry, were recently field isolated [40]. We initially investigated if the uL4(R61D), uL4(R67D), and uL22(K90D) mutant proteins conferred erythromycin antibiotic resistance. We observed that the change uL4(R61D), unlike uL4(R67D), and the uL22(K90D) resulted in resistance to erythromycin (a macrolide) (**Supplemental Figure 1** **and Table 1**). Because uL22(K90D) affects cell growth, we decided to analyze its resistance to a set of antibiotics that target the ribosome or other cellular components by determining the minimal inhibitory concentration (MIC) (**Table 1**). We observed that uL22(K90D) cells had similar sensitivity as wild type cells to most of the tested antibiotics (**Table 1**). However, in addition to erythromycin, uL22(K90D) cells were moderately more resistant to azithromycin (a macrolide) and telithromycin (a ketolide) than the wild type strain by 2- to 4-fold (**Table 1**; **highlighted values**), which are antibiotics whose binding sites are located in the upper segment of the ribosomal exit tunnel [3,41]. These observations indicate that mutations in the non-conserved residues of uL4 and uL22, specifically uL4(R61D) and uL22(K90D) respectively, affect the inhibitory function of antibiotics that target the ribosome exit tunnel. Taken together, these results reveal that uL22(K90D) mutant cells display distinct phenotypic differences from that of wild type and uL4 mutant cells, a slower growth phenotype with a departure from exponential growth at a lower cell density, and an increased resistance to macrolide antibiotic molecules. Our results suggest that the presence of the uL22(K90D) mutation in the ribosome could affect protein synthesis and small molecule binding, leading to a decrease in cell growth and antibiotic resistance. For this reason, we continued our studies using the uL22(K90D) strain.

**Table 1.** Minimum inhibitory concentrations (MIC) of *E. coli* cells expressing wild type uL22 or uL22(K90D) mutant proteins to several antibiotics.

| uL22 | MIC (μg/ml)<sup>1</sup> |  |  |  |  |  |  |  |  |
| --- | --- | --- | --- | --- | --- | --- | --- | --- | --- |
|  | <u>QUI</u> | <u>VIR</u> | <u>CLI</u> | <u>ERY</u> | <u>AZI</u> | <u>TEL</u> | <u>CAM</u> | <u>SPC</u> | <u>AMP</u> |
| **WT** | >512 | >256 | >1024 | **128** | **64** | **64** | 1 | 32 | 4 |
| **K90D** | >512 | >256 | >1024 | **256** | **128** | **256** | 1 | 32 | 4 |

<sup>1</sup> MIC: Minimal Inhibitory Concentration (expressed in μg/mL); QUI: Quinupristin (a streptogramin); VIR: Virginiamycin (a streptogramin); CLI: Clindamycin (a lincosamide); ERY: Erythromycin (a macrolide); AZI: Azithromycin (a macrolide); TEL: Telithromycin (a ketolide); CAM: Chloramphenicol (a phenicol); SPC: Spectinomycin (an aminoglycoside) ; AMP: Ampicillin (cell wall synthesis inhibitor). This table represents the values of at least three independent experiments.

### Global changes in gene expression observed for the uL22(K90D) mutant strain

We next sought to determine differences in gene expression between wild type and uL22(K90D) cells in efforts to understand the observed cell growth behaviors. Ribosome profiling and RNA-seq assays were performed on exponentially grown wild type and uL22(K90D) cells in LB media to quantitate mRNA abundance and translation efficiency in these bacterial strains (**Materials and Methods**). For these assays, cells were harvested at the time point where we started observing the initial growth separation between the cultures (**Figure 2A, arrows**). Comparison of three biological replicates indicated high reproducibility between samples in both normalized RNA and normalized ribosome protected fragments (RPFs) read counts (**Supplemental Figure 2A** **and** **2B**, **respectively**). This reproducibility was also corroborated by a PCA plot with replicates separating according to the tested strains and harvesting dates (**Supplemental Figure 2C**). Plotting RPFs versus mRNA levels indicated good correlation (R^2^=0.66) between both values (**Supplemental Figure 2D**), consistent with the expected differential expression of those genes showing large changes in mRNA abundance and ribosome occupancy. RNA-seq and ribosome profiling data revealed that approximately 8.3% of the 3,729 expressed genes in our uL22(K90D) mutant strain were affected comparing with its parental wild type strain (**Figure 3A**). In the uL22(K90D) strain, 121 genes showed an increase of mRNA levels while 122 genes displayed lower mRNA abundance with respect to the wild type strain; this represents about 78% of the total affected genes. About 15% of the affected genes in the uL22(K90D) strain exhibited changes in their translation efficiency. Specifically, 32 genes showed higher translation efficiency and 22 showed lower translation efficiency in the uL22(K90D) strain compared with wild type. The remaining 7% of affected genes displayed modifications in both mRNA concentrations and translation efficiency. Among these, two genes exhibited higher levels of mRNA together with higher translation efficiency, while ten genes showed low levels of both mRNA and translation efficiency (**Figure 3A**). We did not observe any significant buffered combinations of either low mRNA levels/higher translation efficiency or high mRNA levels/low translation efficiency [42]. Overall, the gene expression data indicate that the uL22(K90D) mutation alters both mRNA abundance and ribosome occupancy on mRNA of certain genes.

**Figure 3.**
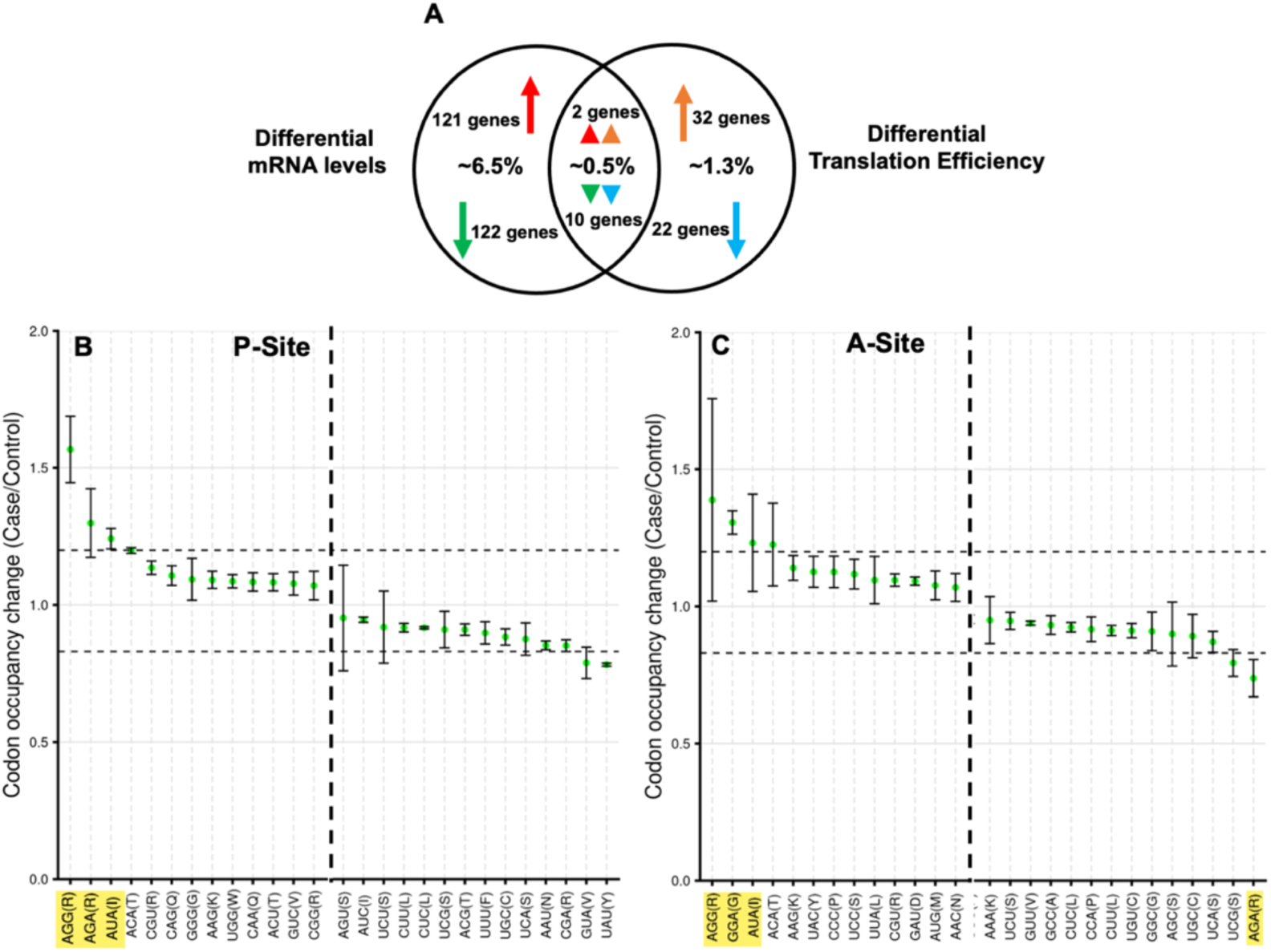
Quantitative changes of gene expression exhibited in the uL22(K90D) mutant strain. (A) Venn diagram representing the distribution of genes showing significantly (log_2_Fold Change>1; adjusted p-value ≤ .05) up- or down-regulation of differentially expressed genes through mRNA levels and/or translation efficiency. Number of genes showing high mRNA levels (red arrows), low mRNA levels (green arrows), high translation efficiency (orange arrows), low translation efficiency (blue arrows), and their combinations are indicated. (B-C) Codon occupancy change of the P-(B), and A-(C), sites observed for the uL22(K90D) strain. Ribosome occupancy values obtained from the uL22(K90D) strain (case) were compared with values from the wild type strain (control). Standard deviations are shown for each codon.

### Ribosome occupancy determined by codon context in the uL22(K90D) strain

Because changes in the ribosomal exit tunnel could affect translation elongation progression [12,43], we decided to compare ribosome occupancy among all 64 codons between the uL22(K90D) and wild type strains (**Materials and Methods**). Our analyses indicated that translating ribosomes in the uL22(K90D) strain have a general tendency to pause on low usage (rare) arginine AGA and AGG codons when occupying the ribosomal P-site (**Figure 3B**). Additionally, the glycine rare codon GGA, as well as AGG codon, were commonly observed at the ribosomal A-site, while arginine AGA codons were avoided in this position (**Figure 3C**).

Rare isoleucine AUA codons also seemed to be frequently observed in both P- and A-site positions (**Figure 3C** **and 3B, respectively**). These data indicate that ribosomes containing uL22(K90D) are more prone to dwelling on rare codons than wild type ribosomes. However, we did not observe any significant enrichment of these rare codons in the genes with low translation efficiency, compared with non-affected genes or with genes with high translation efficiency (**Supplemental Figure 3**). Overall, these data indicate ribosomes containing uL22(K90D) mutant proteins occupy rare codons more often, but the high occupancy does not correlate with codon content of low translation efficiency genes, insinuating that the observed high occupancy is broad among all genes and that genes with low expression might be affected by other translational mechanisms.

### Translationally downregulated genes in the uL22(K90D) mutant strain

Although we did not observe significant enrichments of translationally downregulated genes with related functions, we did observe a pattern in which the first gene of certain functional operons exhibited both lowered mRNA levels and translation efficiency when the rest of the operon displayed only lowered mRNA levels (**Table 2 and Table 3**). Furthermore, these operons contained genes that were primarily related to acid response and carbon metabolism (**Table 2**, **Table 3**, and **Supplemental File**). The most significant change was observed for the *cadB* gene whose translation efficiency was reduced approximately three-fold in the uL22(K90D) mutant with respect to the wild type strain (**Figure 4A** **and Table 2**). The *cadB* gene is part of the *cadBA* operon which is involved in lysine-dependent acid resistance response system in *E. coli* [44]. We observed a significant reduction of ribosomes engaged at the start codon of *cadB* (**Figure 5A**), indicating low translation initiation efficiency. We also observed a significant reduction in the mRNA concentration of both *cadB* and *cadA* (**Figure 4B** **and Table 2**), which suggests low transcriptional expression or high degradation of these mRNA sequences. The expression of genes within this operon is controlled by several transcription factors, including GadE-RcsB and CadC [45–47]. We did not observe reduction in the expression of the *rcsB* gene (**Supplemental File**), nor the *gadE* gene; rather, this last regulatory gene shows high mRNA levels as discussed below (**Figure 4B** **and Table 2**). In the case of *cadC*, despite the lack of significant changes in mRNA levels and translation efficiency (**Supplemental File**), we observed low ribosome protection density at the *cadC* 3’-end sequences after its ^120^PPP^122^ sequence (**Figure 5B**), which is a ribosome arresting sequence studied previously [48]. We suspect that the translational expression of *cadC* is stalled at the ^120^PPP^122^ sequence in the uL22(K90D) strain, reducing CadC protein expression, and thus could reduce the transcription of the *cadBA* operon as observed in our data (**Figure 4A** **and 4B, and Table 2**). The gene *adiY* also displayed reduced translation efficiency in the uL22(K90D) strain. Like *cadB*, *adiY* is also part of the acid response and activates the expression of *gadA* and the *gadBC* operon [49]. Similar to what was observed in *cadB*, the *adiY* transcript showed a reduction of ribosome occupancy at its start codon (**Figure 5C**). These data indicate that the expression of *adiY*, like *cadB*, could be affected during translation initiation as well. Overall, in the uL22(K90D) strain, we observed the reduction of translation efficiency in genes involved in various acid response mechanisms. The reduction in the expression of these genes seems to correspond with an increase in the expression of acid and alkaline resistance genes, especially those consisting of the *gad* regulon system (**Figure 4B** **and Table 2**) with *cadBA* being the only genes in the regulon that are downregulated.

**Figure 4.**
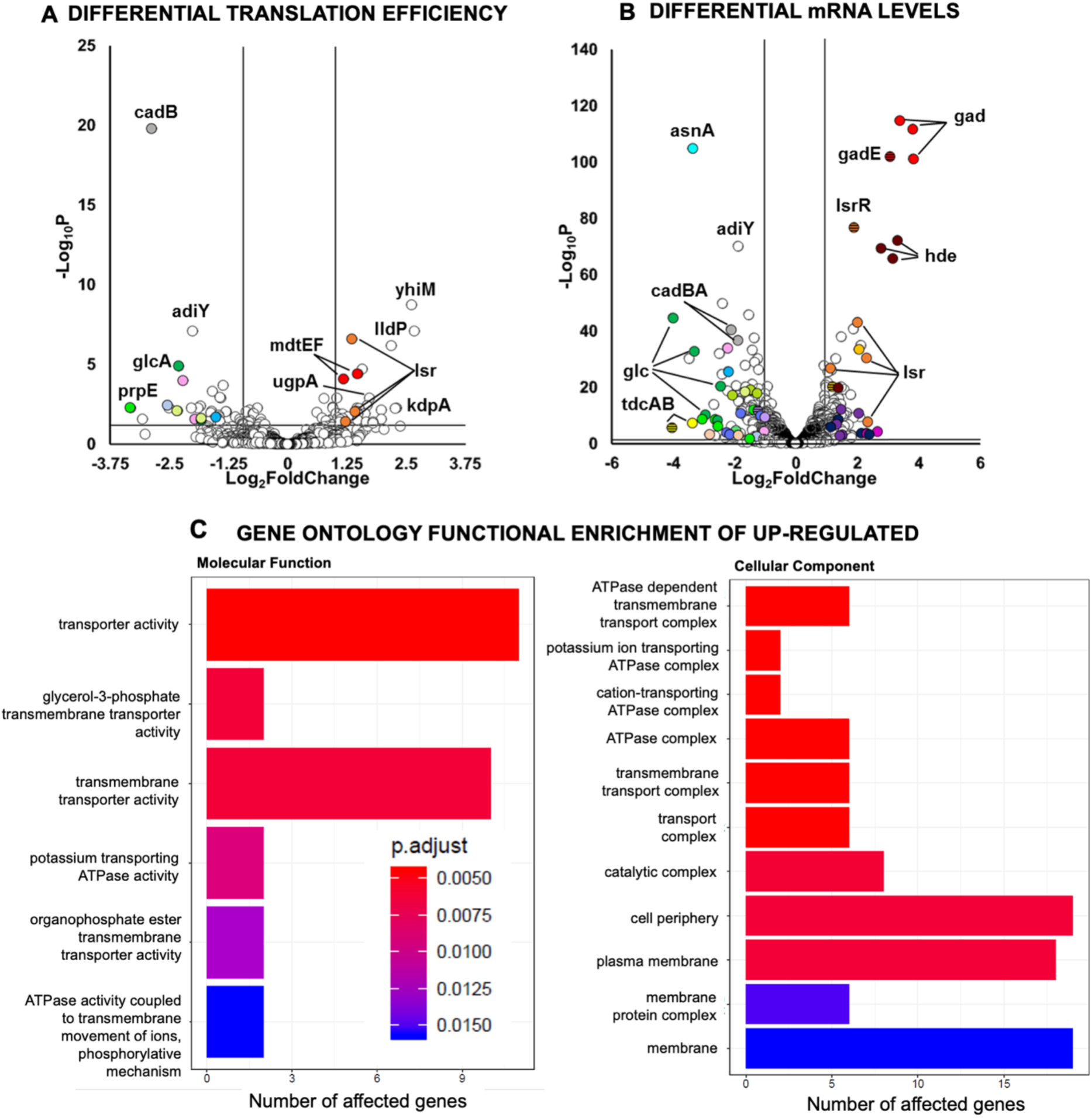
Functional groups and genes which expression is affected in the uL22(K90D) mutant strain. (A) Volcano plot showing significance vs changes in translation efficiency of all tested genes. (B) Volcano plot showing significance vs changes in mRNA levels. (C) Graphical representations of statistically significant enriched GO terms, molecular function and cellular components observed in the mutant strain. All genes >1 log_2_ fold change with an adjusted p-value ≤ .05 show significant changes, and outlying genes and operons discussed in this study are labeled. The p. adjust heathmap applies to both panels. This data represents the analysis of three biological replicates.

**Figure 5.**
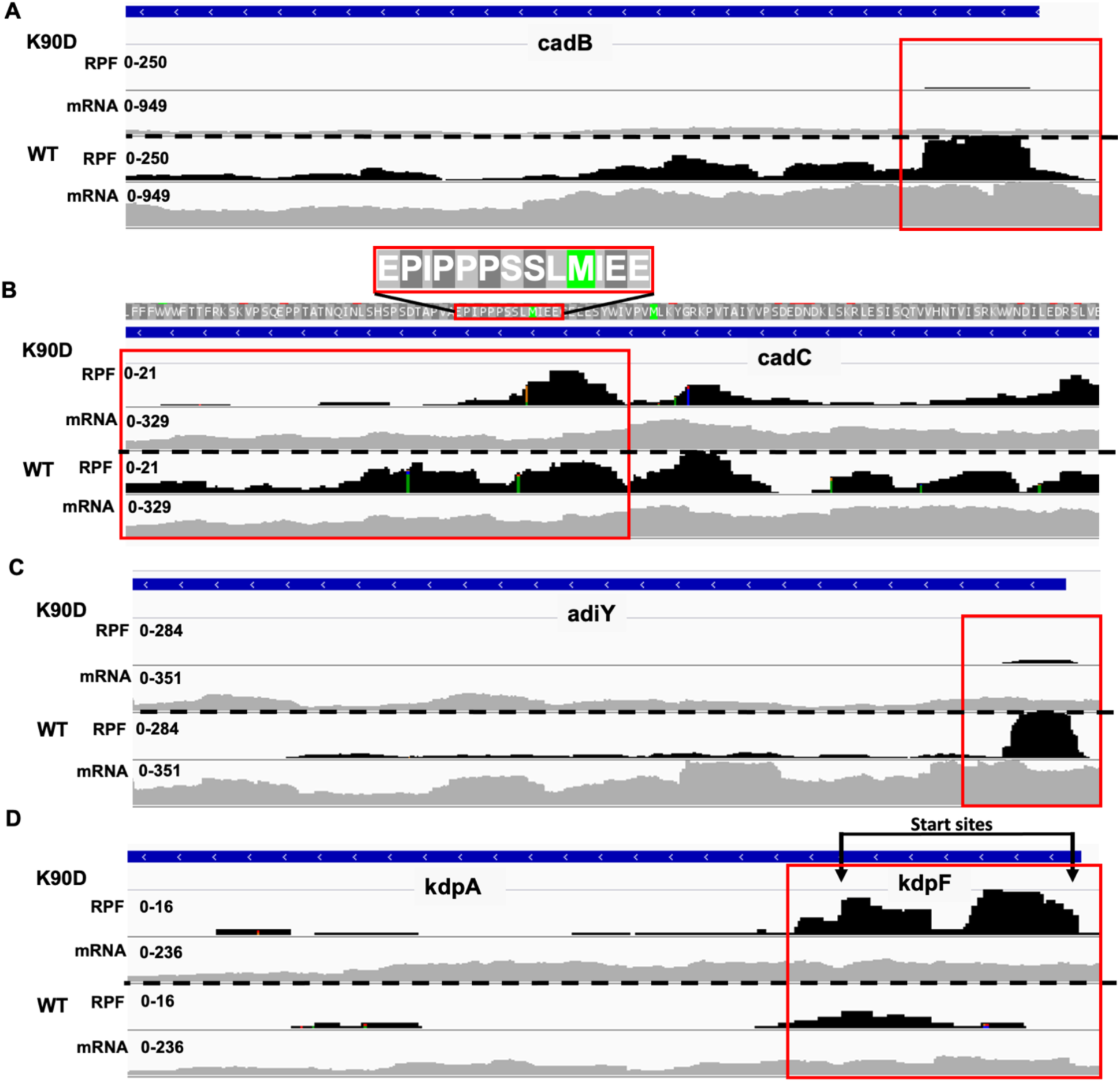
RPF and mRNA coverage profiles of affected genes in the uL22(K90D) mutant strain. Individual gene coverage profiles visualized with IGV of RPF (black) and mRNA (gray) for uL22(K90D) (top panels) and wild type (bottom panels) samples. Each panel is auto scaled for each gene and by group. RPF and mRNA read count units are arbitrary. (A) *cadB*, (B) *cadC*, (C) *adiY*, and (D) *kdpFA.* Differences between translation for each gene are highlighted in red boxes. Each track is a merged (summed) representation of the data from three independent experiments.

**Table 2.**
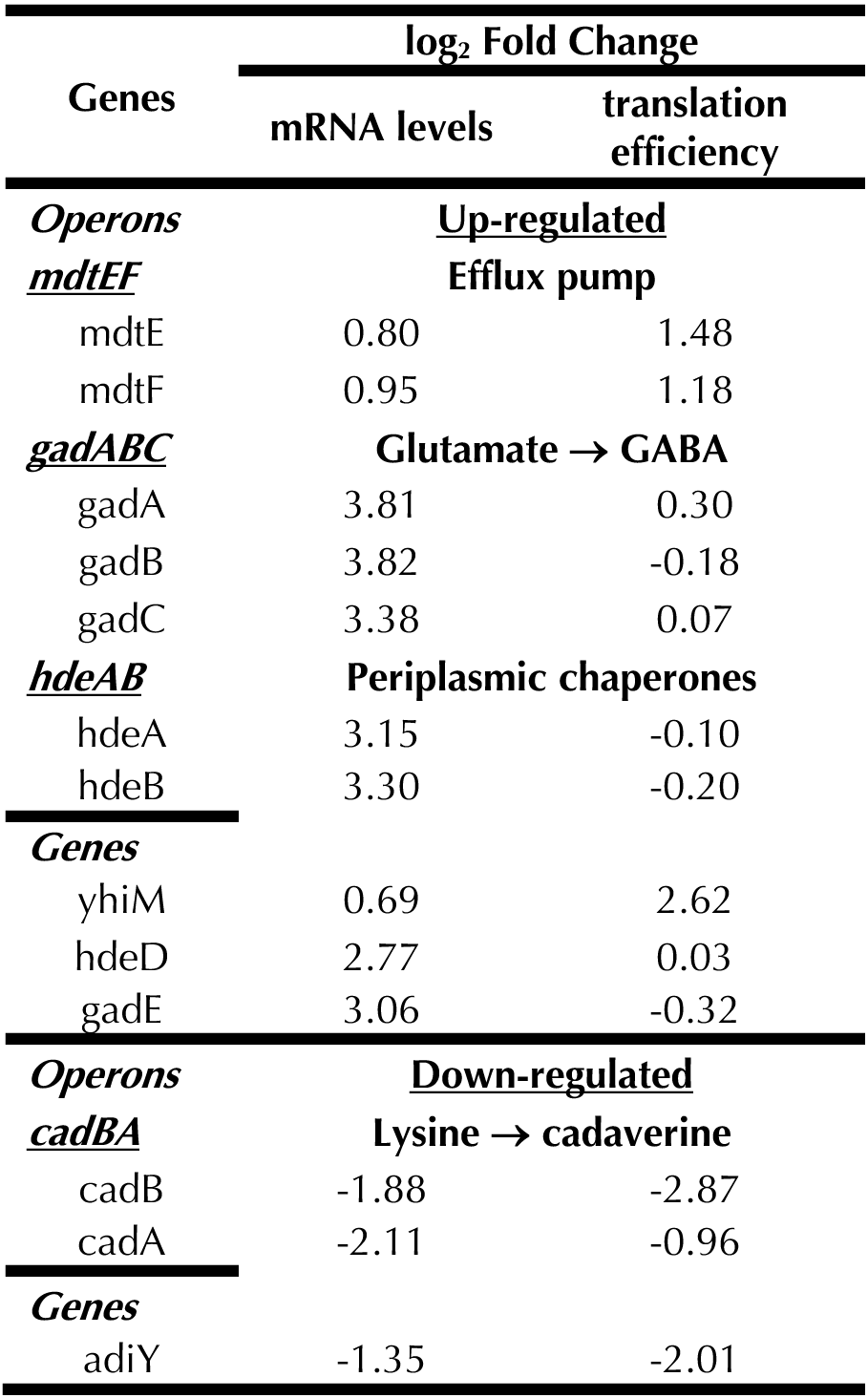
Acid resistance genes affected in the uL22(K90D) strain

**Table 3.**
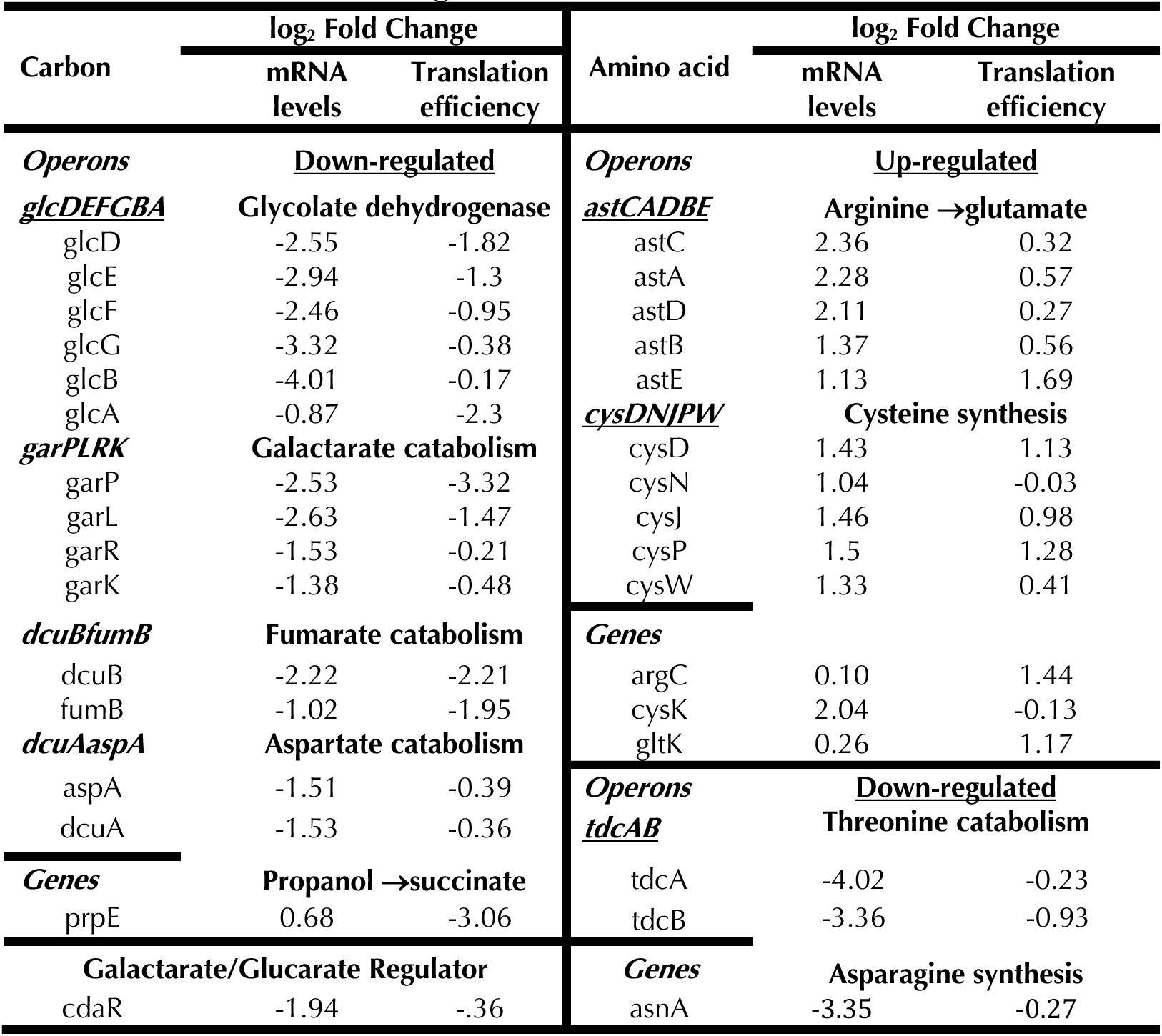
Metabolic related genes affected in the uL22(K90D) strain

### Metabolically related genes affected in the uL22(K90D) strain

We observed changes, mostly reductions, of mRNA levels and translation efficiency of genes related to alternative carbohydrates catabolism in the uL22(K90D) mutant strain [50,51]. Operons involved in the catabolism of glucarate, galactarate, glycerate, glycolate, and glyoxylate degradation all showed low mRNA levels (**Table 3**). The transcriptional regulator *cdaR* gene [52] that controls the glucarate and galactarate operons exhibited low mRNA levels as well (**Table 3**). However, the regulator of the glycolate degradation operon, *glcC*, did not show any significant changes in both mRNA levels or translation efficiency (**Supplemental File**).

Interestingly, as seen for the *cadBA* operon, the first gene of these affected operons exhibited significant reductions of their translation efficiency in the uL22(K90D) mutant strain (**Table 3**), which could imply a decoupling effect of transcription from translation promoted by a slow translation of the leader gene of the operon. In concordance with changes in using alternative sources of energy, we observed that genes related to the electron transport chain and redox under anaerobiosis contributed to the altered gene profile in uL22(K90D) mutant cells. Several operons known to be associated with the stress response and electron transport chain components contain genes with a pattern of significantly altered mRNA levels in the uL22(K90D) cells compared to the wild type (**Table 4**). In general, reductase activities were downregulated. One of these operons is the fumarate reductase operon (**Table 4**), whose substrate is produced by the alternative carbon activities whose expression is reduced as well (**Table 3**) [53]. This suggests that metabolism of fumarate could be altered. Alternatively, complexes involved in oxidoreduction of other metabolites, such as succinate, were upregulated (**Table 4**), perhaps compensating the reduction of these pathways that utilize fumarate as an intermediate metabolite. Our results suggest that the pattern of differential up/down alterations in gene expression and translation could be attributed to favoring certain catabolic and oxidative stress mechanisms.

**Table 4.**
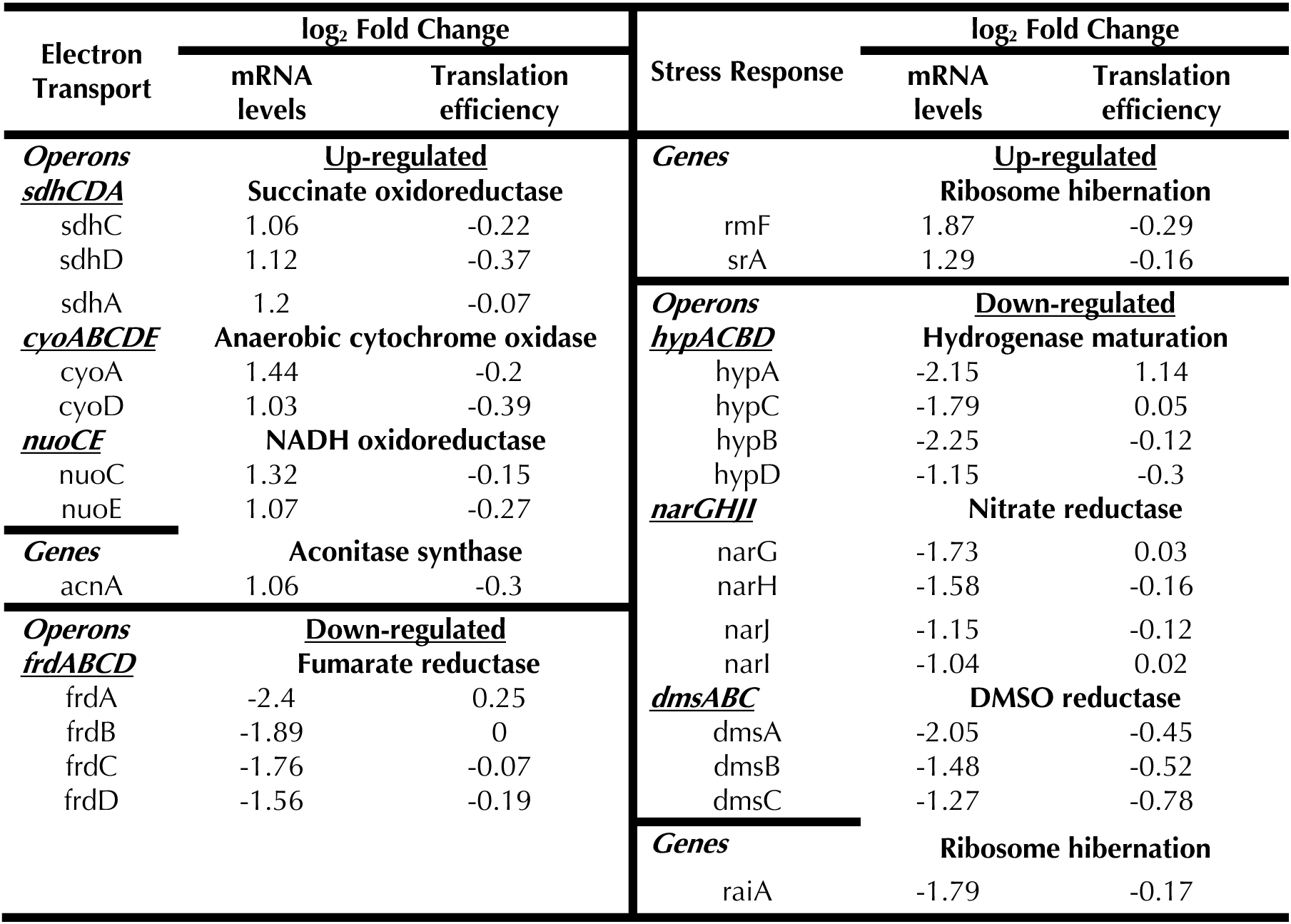
Electron transport and stress response related genes affected in the uL22(K90D) strain

Not only does the uL22(K90D) mutant appear to have altered alternative carbohydrate metabolism, it also appears to have alterations in amino acid metabolism. Several operons involved in amino acid metabolism showed downregulation and we noticed that changes in these pathways are mostly related with the consumption and generation of ammonia. As expected, the *tna* operon showed low mRNA levels (**Table 5**) and we confirmed that production of the tryptophanase enzyme was reduced in the uL22(K90D) mutant strain as well (**Supplemental Figure 4**). This indicates reduced degradation of available L-Trp and L-Cysteine, as well as a decrease of indole, hydrogen sulfide, ammonia, and pyruvate production [12]. Besides the *tna* operon, the *asnA* gene exhibited a significant reduction in its mRNA levels as well (**Table 3**).

**Table 5.**
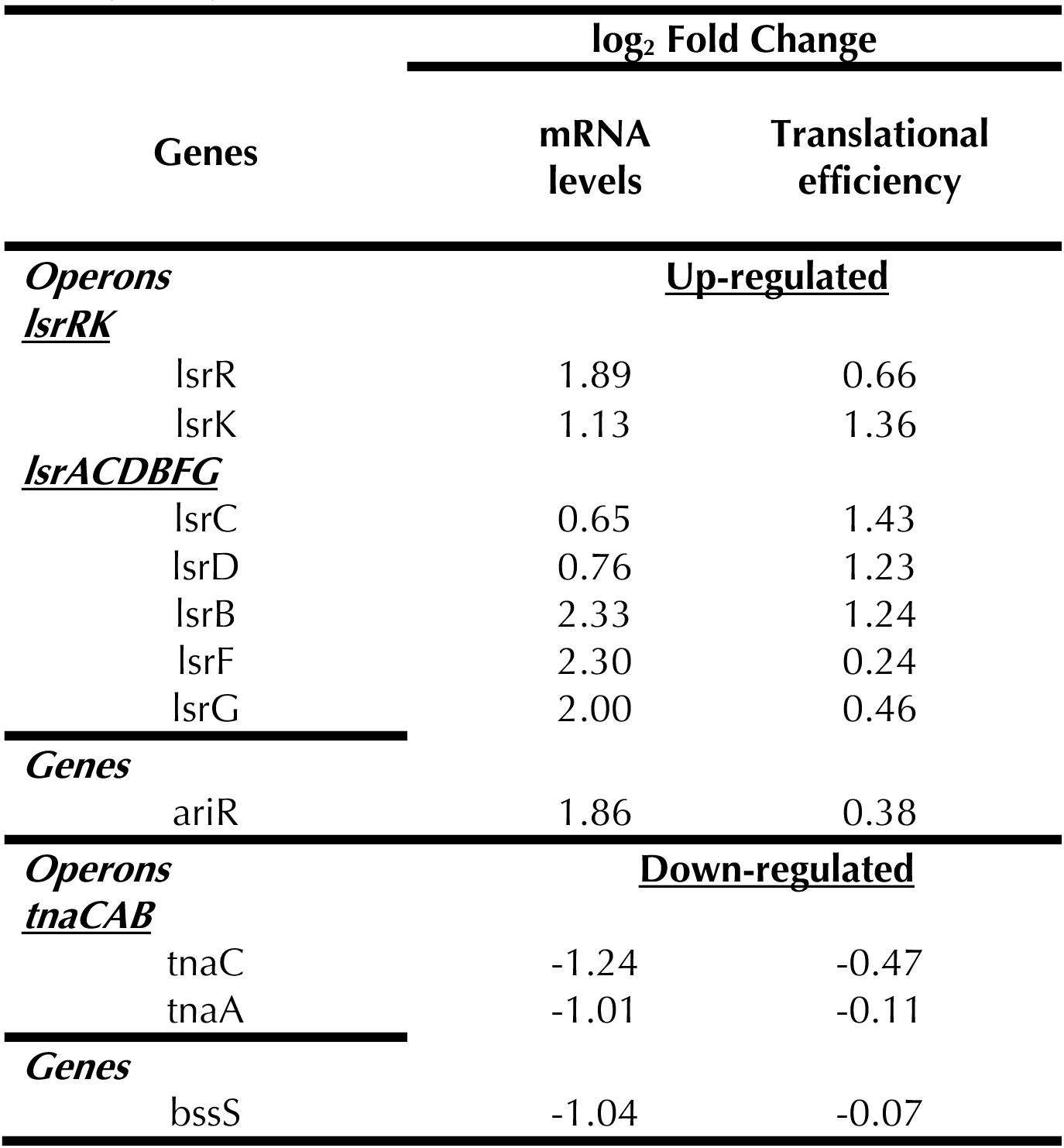
Biofilm formation related genes affected in the uL22(K90D) strain

*asnA* is one of two asparagine synthetases in *E. coli* that converts aspartate into asparagine, consuming ammonia and increasing inorganic phosphate and protons as byproducts [54]. Similarly, the *tdcA* and *tdcB* genes of the *tdcABCDEFG* operon involved in L-threonine and L-serine catabolism were significantly downregulated (**Table 3**). Activity of this operon generates ammonia, pyruvate, and propionyl-CoA as byproducts during anaerobiosis [55]. Contrary to those repressed operons, the alkaline resistance *astCADBE* operon which catabolizes arginine, producing ammonia and generating glutamate [56], exhibited increased mRNA levels (**Table 3**). Interestingly, *argC*, which is involved in the arginine biosynthetic pathway, also showed increased translation efficiency in the uL22(K90D) mutant strain (**Table 3**), perhaps contributing to the generation of glutamate by funneling arginine. Furthermore, operons involved in the metabolism of sulfur, alkaline resistance, and the synthesis of cysteine (*cysB*, *cysDNC*, *cysJIH*, *cysK*, *cysPUWAM*) [57] showed significant increases in their mRNA levels (**Table 3**). Thus, in general, we observed a tendency toward changes in amino acid metabolism that would modify the flux of pyruvate, ammonia, sulfur, and protons.

### Translationally upregulated genes in the uL22(K90D) mutant strain

Gene Ontology (GO) and Kyoto Encyclopedia of Genes and Genomes (KEGG) enrichment analysis for genes with differential translation efficiency in the uL22(K90D) mutant strain revealed significant enrichment in the up-regulation of transporters, transporter activity, cell periphery, and plasma membrane related genes (**Figure 4C; Supplemental File**). The genes, *ugpA*, *kdpA*, *mdtE*, and *mdtF* constituting three operons that express protein subunits of transporters, together with the *yhiM* and *lldP* gene transporters, showed high levels of translation efficiency (**Figure 4A**). The remaining genes that constitute the *kdpFABC* and *ugpBAEC* operons, which express transporters that carry potassium and phosphate into the cells respectively [58, 59], exhibited high mRNA levels (**Supplemental File**). We noticed that the *kdpF* gene (31 codons long) and the *kdpA* gene, located at the front of the *kdpFABC* operon, showed two significant ribosome protection signatures in the uL22(K90D) samples comparing with the wild type samples (**Figure 5D**). Both ribosome protections correspond to the translational start sites of *kdpF* and *kdpA*, which indicates that translation initiation of these two genes is enhanced in the uL22(K90D) mutant strain. The *mdtEF* operon, a multidrug efflux transporter, also showed high translation efficiency. High expression of the *mdtEF* operon was accompanied by high mRNA concentrations of the transcriptional regulator *gadE* gene (**Figure 4B** **and Table 2**), which is located upstream of the *mdtEF* operon. These observations suggest that there might be high expressional activity of *arrS-gadE-gadF-mdtEF* chromosomal loci [60] in the uL22(K90D) strain. *gadE* and the regulatory ncRNAs control the expression of genes that are involved in conferring cellular resistance to acid induced stress [60,61]. In agreement, we observed high mRNA levels of acid response genes and operons, among them the *hde* and *gad* operons (**Figure 4B** **and Table 2**) and the membrane protein *yhiM*, which also showed increased translation efficiency (**Figure 4A**; **Table 2**). This final set of data indicate that the uL22(K90D) strain highly expresses acid response genes. Overall, these results reveal that in the uL22(K90D) strain there is an increase in membrane transportation activity accompanied by an increase in glutamate-related acid resistance activities.

Because a significant number of genes affected in the uL22(K90D) strain are associated with the acid and alkaline stress response systems, we sought to investigate if these changes had a functional meaning in the growth of this uL22(K90D) mutant strain. This was accomplished by challenging the uL22(K90D) strain to grow under highly acidic and alkaline conditions. We reasoned that the differentially expressed genes related to acid and alkaline resistance would provide growth advantage of the uL22(K90D) over its parental strain in acid and alkaline growth conditions compared to growth in neutral conditions. We evaluated the difference in cellular density between wild type and uL22(K90D) cultures during exponential growth in media with varying pHs, ranging from 4.6, 7.0, to 9.2 (**Figure 6A**). We observed that, at comparable cellular densities, the differences in growth between wild type and uL22(K90D) were much smaller in both acidic and basic conditions compared with neutral growth conditions (**Figure 6B**). These results imply that uL22(K90D) strain is more competitive with the wild type strain when grown in acidic and basic conditions. However, since the rate of exponential growth in uL22(K90D) cultures is still lower than that of wild type cultures in all tested conditions, this indicates that there are additional genes, independent of acidic and basic stress responses, that negatively affect the growth of the uL22(K90D) strain in rich media. These genes could be those involved in metabolism as indicated above.

**Figure 6.**
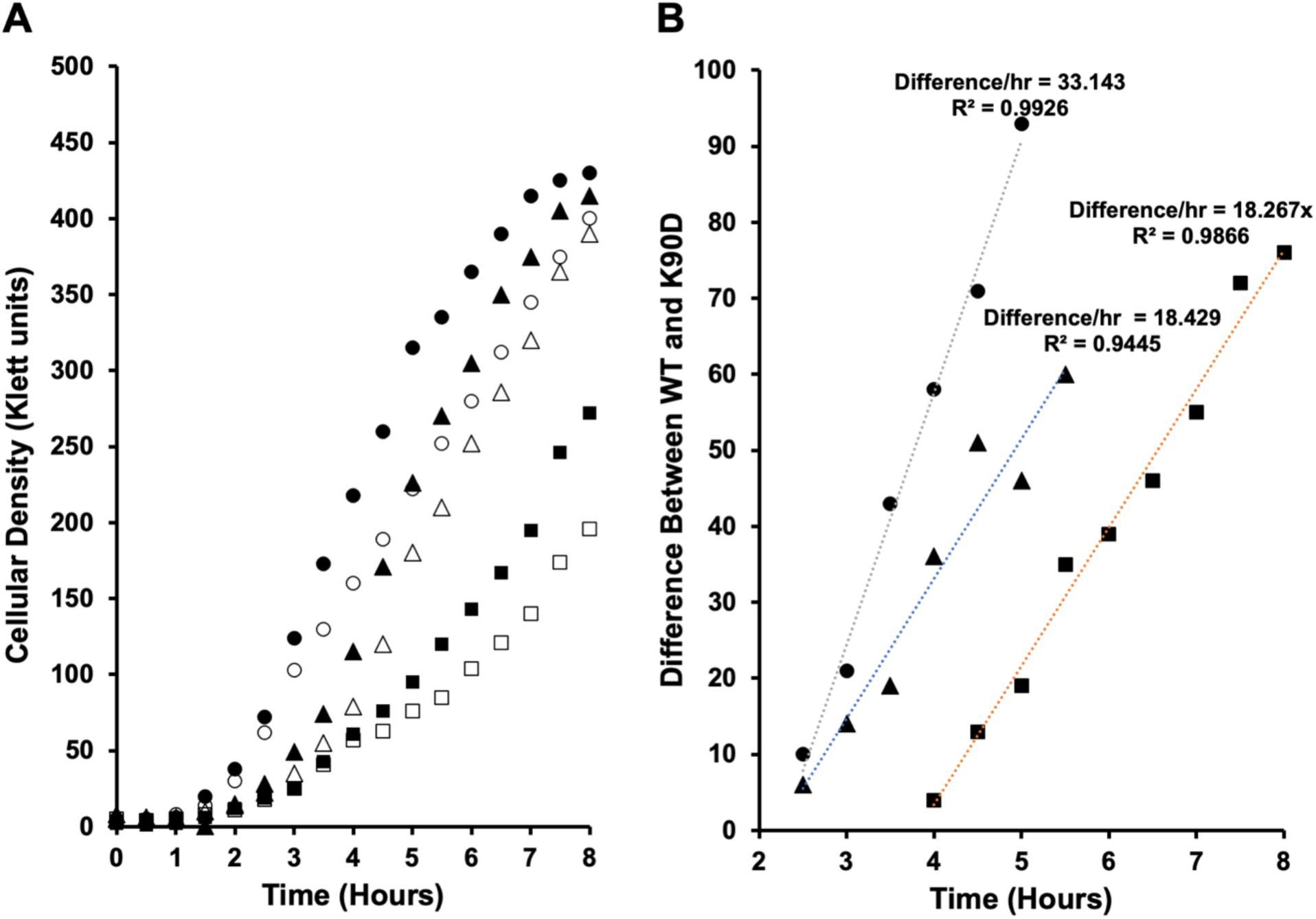
Growth of wild type and uL22(K90D) strains in acidic and basic conditions. (A) Plot showing growth curves of wild type (closed) and uL22(K90D) (open) cultures performed in LB media with a pH of 4.6 (triangles), 7 (circles), or 9.2 (squares). B) Difference in cellular density between wild type and uL22(K90D) cultures during exponential growth phase observed in A) were plotted versus time. Data shown are representative of three biological replicates.

### Affected genes involved in biofilm formation

Our observations indicate that uL22(K90D) mutant cells reach the stationary phase at a reduced cellular density compared to the wild type strain (**Figure 2A**), which could imply changes in quorum sensing. The synthesis and degradation of two important interspecies signaling molecules, indole and AI-2, seem to be affected in our mutant strain. As indicated above, indole production in uL22(K90D) mutant cultures was reduced. Additionally, we observed an increase in mRNA levels of the operons involved in the uptake and phosphorylation of the AI-2 molecule, *lsrACDBFG* and *lsrRK* (**Figure 4B** **and Table 5**). Interestingly, *lsrK* also showed a significant increase in its translation efficiency (**Figure 4A** **and Table 5**). Both indole and AI-2 are known to modulate the expression of other genes, including genes related to the pH response, quorum sensing, and biofilm formation [62,63]. The expression of several genes activities that link indole and AI-2 were also affected in the uL22(K90D) strain. The gene *bssS*, which responds to glucose and AI-2, reducing biofilm formation and increasing the uptake of indole [64], exhibited low mRNA levels in the uL22(K90D) mutant strain compared to wild type (**Table 5**).

Conversely, the *ariR* gene, a biofilm regulator linked to indole, AI-2, and acid resistance [65], showed a significant increase in mRNA levels in the uL22(K90D) mutant (**Table 5**). These observations indicate that the uL22(K90D) change affects the expression of genes involved in the metabolism and functionality of cell signals related with biofilm and quorum sensing. We investigated whether these changes in expression led to alterations in the formation of biofilms in cultures of the uL22(K90D) strain. Results using a standard biofilm crystal violet assay (**Materials and Methods**) revealed that uL22(K90D) cells form biofilms more readily, with an average increase of 42%, compared to wild type cells (**Figure 7**).

**Figure 7.**
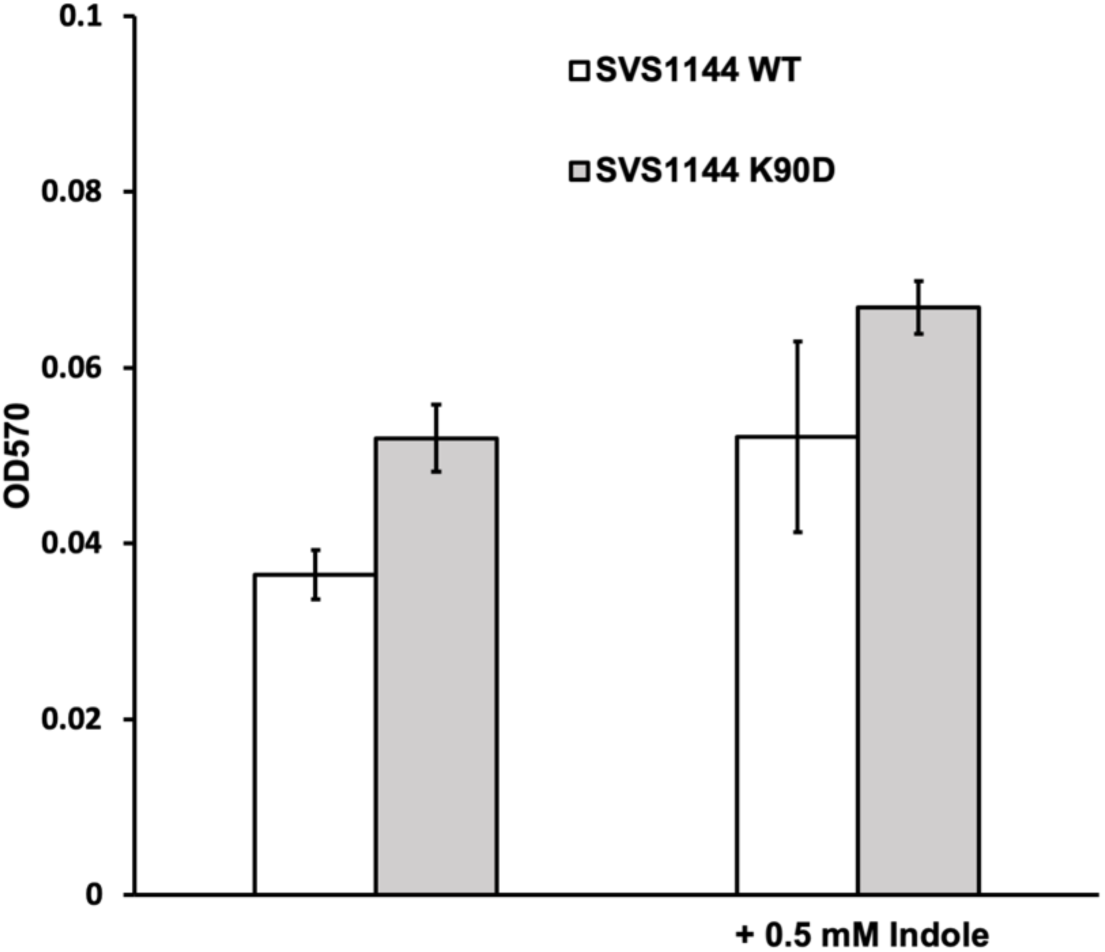
Biofilm formation observed in wild type and uL22(K90D) strains. Biofilm formation was tested in LB media in the absence (n=6) or presence of 0.5 mM indole prior to 24 hours of static incubation (n=2) as indicated in **Materials and Methods**. Error bars are based on standard deviation.

Since indole has been implicated as a biofilm inhibitor in most *E. coli* strains, we were interested to see if the increased biofilm formation observed for the uL22(K90D) cultures was due to its defect in indole synthesis. If so, we would expect that the addition of indole would decrease biofilm formation of uL22(K90D) cultures to that observed in wild type cultures. However, we observed that biofilm formation increased for both the wild type and uL22(K90D) cultures when supplemented with indole and there was still a disparity present in the generation of biofilms between the uL22(K90D) and wild type cultures (**Figure 7**). This suggests that the increase in biofilm formation observed for the uL22(K90D) cells is not due to its reduction of indole synthesis. Supporting this observation, we observed negligible differences in the formation of biofilms between an isogenic indole mutant, CY15000 (**Supplemental Table I**), and the same strain complemented with the *tnaA* gene (**Supplemental Figure 5**). These observations indicate that the decreased indole production is not the definite factor that influences the increase of biofilm formation in our uL22(K90D) strain.

## DISCUSSION

In this work, we demonstrated that a single mutational change at a non-conserved residue position of the uL22 protein significantly impacts cellular growth and gene expression. This mutational change added an acidic amino acid in a position where only basic or neutral amino acids residues are selected among prokaryotes (**Figure 1B**). Cells expressing ribosomes with the uL22(K90D) mutant protein showed growth deficiencies in rich media (**Figure 2**) and were resistant to a set of macrolide antibiotics (**Table 1**). Changes in the expression of several genes correlated with the observed cell growth phenotype. 123 genes show high levels of mRNA steady state concentrations, while a comparable number of 132 was observed for genes showing low levels of mRNA (**Figure 3A**). Similarly, 34 genes showed an increase in translation efficiency, while 32 exhibited low translation efficiency (**Figure 3A**). Translational traffic under ribosomes with uL22(K90D) mutant proteins displayed a shift of ribosomes that spend longer time at rare arginine, isoleucine, and glycine codons (**Figure 3B-C**). We noticed that genes with low translation efficiency in the uL22(K90D) strains are mostly located at the front of several operons, where downstream genes of the operon exhibited low mRNA levels (**Table 3**). Genes such as *cadB* and *adiY*, both related to acid response, were the most significantly translationally downregulated (**Figure 4A** **and Table 2**). mRNAs of both genes displayed low ribosome occupancy at their start codons, indicating reduced translation initiation (**Figure 5A** **and 5C**). Additionally, the *cadC* transcriptional regulatory gene, which controls the expression of the *cadBA* operon, exhibited low ribosome protection signals after a translation arrest site located in a region rich in glutamic and proline codons (**Figure 5B**). This indicates that this gene may not be expressed, which correlated with low mRNA levels of the *cadBA* operon (**Figure 4B**). The expression of other genes related to the catabolism of alternative carbon sources (glucarate, galactarate, glycerate, glycolate and glyoxylate) and amino acid catabolism (*tnaA*, *asnA* and *tdcAB*) were also down-regulated (**Figure 4A**, **4B, and Table 3**), together with a reduction of genes involved in the utilization of alternative oxidizing agents used during anaerobiosis (**Table 4**). On the other hand, upregulated expression was observed for genes and operons expressing transporters and membrane components (**Figure 4C**), genes related to the acid and alkaline response (**Table 2**), the *lsrR* regulon involved in the uptake and phosphorylation of AI-2, and the *ariR* gene that controls both biofilm formation and the acid response (**Table 5**). These upregulated functions seem to produce an enhancement in acid and alkaline resistance (**Figure 6**) and in the production of biofilms (**Figure 7**). In summary, our data indicate that the uL22(K90D) change in the uL22 protein reduces the expression of genes, which induces compensatory increases in the expression of genes with related activities, thus altering cellular behaviors.

Comparing our results with previous analyses of bacteria resistant to macrolides, we found a considerable number of differences and commonalities. In the case of the uL22(Δ82-84) mutational change, the regulation of the *secMA* operon, whose function is important for protein translocation through the cellular membrane [66], is affected [17]. In the case of the uL22(K90D) mutational change, we did not observe any significant changes in the expression of the *secMA* operon (**Supplemental File**). Both uL22(Δ82-84) and uL22(K90D) mutations produced a reduction in *tnaA* mRNA abundance, albeit the 2-fold reduction (**Table 4**) in our uL22(K90D) mutant was far more modest than the 67-fold reduction in the uL22(Δ82-84) [17]. These observations led us to conclude that different mutational changes in the uL22 protein generate different outcomes regarding the global expression of genes, suggesting that variability in the nature of the tunnel could have different effects regarding the gene being translated, as previously suggested [6].

### Translational effects produced by the uL22(K90D) mutant protein

Ribosomes with uL22(K90D) mutant proteins are prone to stay longer at rare codons (**Figure 3B-C**), especially arginine codons. Also, arginine catabolism seems to be increased to generate glutamate (**Figure 8**) which would reduce the pool of available arginine to produce arginyl-tRNA and enhance the ribosome occupancy at arginine rare codons. We suspect that these effects generate translational stress, which induces the expression of the ribosome hibernating factors RmF and SrA (**Table 4**) and reduces expression of the RmF-competitor, RaiA factor (**Table 4**) [67]. Because ribosomes are being sequestered, this could reduce translation initiation of genes dependent on ribosome concentrations, including *cadB, adiY*, and *prpE*, as well genes in the front the *glc*, *gar*, and *dcuB-fumB* operons (**Figure 8**). It is quite possible that reduction in the expression of all of these genes could be the main factor that produces the delay in cell growth observed in the mutant strain (**Figure 2**).

**Figure 8.**
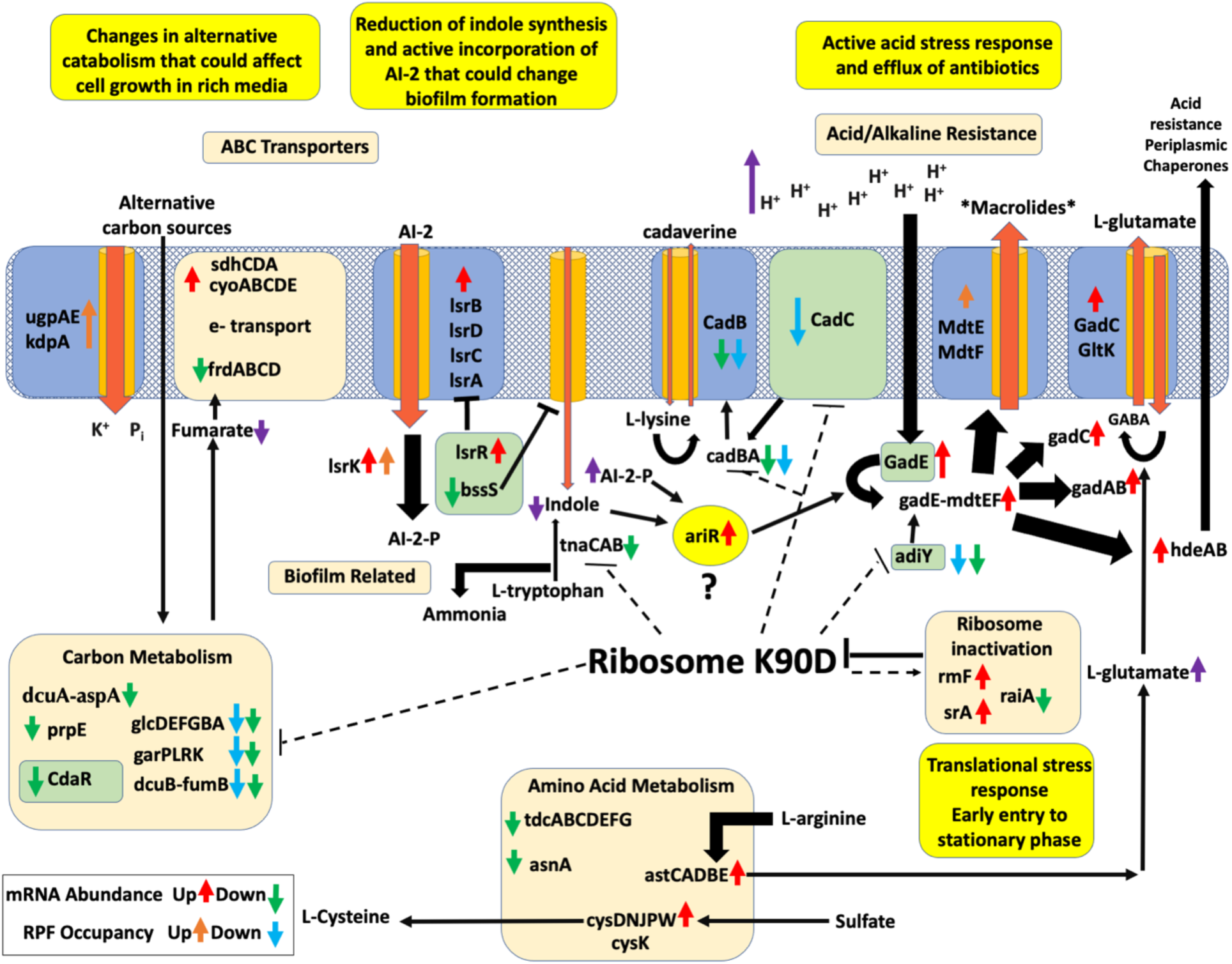
Model of gene expression regulation observed in the uL22(K90D) mutant strain. This figure represents the interconnections between genes and substrates affected in the uL22(K90D) strain. Dotted lines mark genes showing reduced translation efficiency due to the activity of ribosomes containing uL22(K90D) mutant proteins. The gray mesh box represents the inner cellular membrane of *E. coli*. Blue boxes with orange cylinders, or just orange cylinders, represent membrane channels and transporters. Orange arrows indicate movement of molecules across the channels and transporters. Green boxes enclose regulatory genes. Yellow circle encloses a possible key gene that controls the phenotypes observed in the uL22(K90D) mutant strain. Black arrows indicate movement of metabolites, chemical reactions, or proteins interactions with targeted genes, the thickness of the arrow is proportional to the activity of the event. Magenta arrows represent the level of accumulation expected for certain metabolites.

Mutant ribosomes also exhibit arrest at certain sequences. As seen for the *cadC* gene, ribosome coverage at its 3’-end is notably low following the PPP stalling motif (**Figure 5B**), which suggests that the CadC protein regulator may not be produced. This stalling motif is preceded by a rich region of glutamic acid residues, located between 6-11 residue positions into the exit tunnel, precisely in the upper region of the tunnel before reaching the constriction region. Context of the upstream sequence is important for PPP motif stalling [70]; it could be possible that the presence of the aspartic acid residue in the 90th position of uL22 could affect the movement of this nascent peptide through the constriction region, increasing the timing of the ribosomal arrest at the PPP motif.

### Acquisition of antibiotic resistance

It has been shown in previous studies that high activities of efflux pumps contribute greatly to uL22 antibiotic resistance mutations [71,72], whereas elimination of the expression of core metabolic genes also contributes to antibiotic resistance [73]. The genes *mdtE* and *mdtF*, part of the *MdtEF-TolC* efflux pump system, are expressed at a higher level in the uL22(K90D) mutant, more significantly through a translational effect than mRNA levels (**Figure 4A**; **Table 2**). Also, in our study, *prpE* (propionyl-CoA synthetase) was significantly translationally downregulated in the uL22(K90D) mutant compared to wildtype (**Figure 4A**; **Supplemental File**). It is likely that the upregulation of the *mdtEF* efflux pump genes and the reduction in the expression of *prpE* could contribute to the erythromycin resistant phenotype observed in the uL22(K90D) mutant. Our data indicate that the high transcriptional activity of *gadE* and surrounding genes could explain the increase of the *mdtEF* efflux pump genes, and as a consequence the resistance to erythromycin, as previously seen [74]. However, we cannot eliminate the possibility that this mutation could also affect erythromycin binding and/or movement into the ribosomal exit tunnel, as we expect to be the for case for the observed telithromycin resistance based on the structure (**Table 1**) [3].

### Biofilm formation in the uL22(K90D) strain

The uL22(K90D) strain has an increased capacity to generate biofilms and sediment, compared to its parental strain (**Figure 7**; **Supplemental Figure 6**). The two signaling molecules, indole and AI-2, have been implicated in the formation of biofilms in *E. coli* [75,76]. However, indole and AI-2 seem to play opposite roles in *E. coli*. Indole is most commonly associated with inhibition of biofilm formation [76] while AI-2 stimulates biofilm formation [75]. Also, indole has been implicated in cellular repulsion [76] while AI-2 is involved in cellular attraction [75]. Furthermore, indole is secreted by cells upon transition into the stationary phase, which corresponds to the timing in which AI-2 uptake by cells occurs [77,78]. Production of indole is solely controlled by the *tna* operon, whose expression is downregulated through mRNA levels in uL22(K90D) cells (**Table 5**). Consistent with changes in the expression of *tnaA*, we observed a dramatic reduction in the synthesis of tryptophanase (**Supplemental Figure 4**) which leads to lower indole production. In our experiments, the addition of indole to both wild type and uL22(K90D) cultures increased the formation of biofilms (**Figure 7**). However, our data shows a reduction of the *bssS* gene (**Table 5**) that it is known to induce the incorporation of indole into the cell [64]. All these changes suggest a reduction of indole within the uL22(K90D) cell (**Figure 8**). On the other hand, genes related to AI-2 uptake and phosphorylation were highly upregulated in the uL22(K90D) strain (**Table 5**), indicating a reduction of AI-2 extracellular activity. We suspect that incorporation and phosphorylation of AI-2, together with reduction of internal indole, may explain the increase in biofilm formation and sedimentation phenotypes observed in uL22(K90D). Supporting this idea, we observed an increase in the expression of the *ariR* gene (**Figure 8**), that not only increases the formation of biofilms, but also increases the expression of the *gad* operon (**Figure 8**) [65]. We suspect that the *ariR* gene could be a key regulator of both biofilm and acid response/resistance to macrolides phenotypes in our mutant strain.

In conclusion, our results indicate that a single amino acid change in the extended loop of the uL22 protein has significant effects in the expression of genes related to the acid resistance response, metabolism, cellular indole concentrations, and the incorporation and modification of AI-2. These changes are accompanied by a deficiency in cell growth, an enhancement in biofilm formation, and an increased resistance to particular macrolides and acidic conditions. Current work should focus on determining the molecular mechanism by which the uL22(K90D) mutant is modulating changes in gene expression and translation of the discussed genes.

## DATA AVAILABILITY

Ribosome profiling data was analyzed using RiboToolkit, a free webtool available at (http://rnabioinfor.tch.harvard.edu/RiboToolkit/). Data was also prepared for visualization on a genome viewer using RiboGalaxy, a free webtool available at (http://ribogalaxy.ucc.ie).

## ACCESSION NUMBERS

The Ribo-seq and RNA-seq data reported in this paper have been deposited in the NCBI Gene Expression Omnibus (GEO) database with accession number <u>GSE171533.</u>

## SUPPLEMENTARY DATA

Supporting File

## ACKNOWLEDGEMENTS

The authors would like to thank Kevin D. Young for supplying the plasmids pLP8 and pTnaA; Sean Moore for the plasmid pS10, phage P1vir, and the cell line SM1110 ; and Nora Vazquez-Laslop for the antibiotics quinupristin, virginiamycin S1, azithromycin, telithromycin, clindamycin. They also would like to thank Elizabeth Ramsey and D. Emma Bryant for their work collecting data. LRCV would like to thank the UAH provost office, college of science dean, and Paul Wolf chair of the biology department for their financial support used in the preparation of the ribosome profiling assays.

## SUPPLEMENTARY DATA

**Supplemental Table1.**
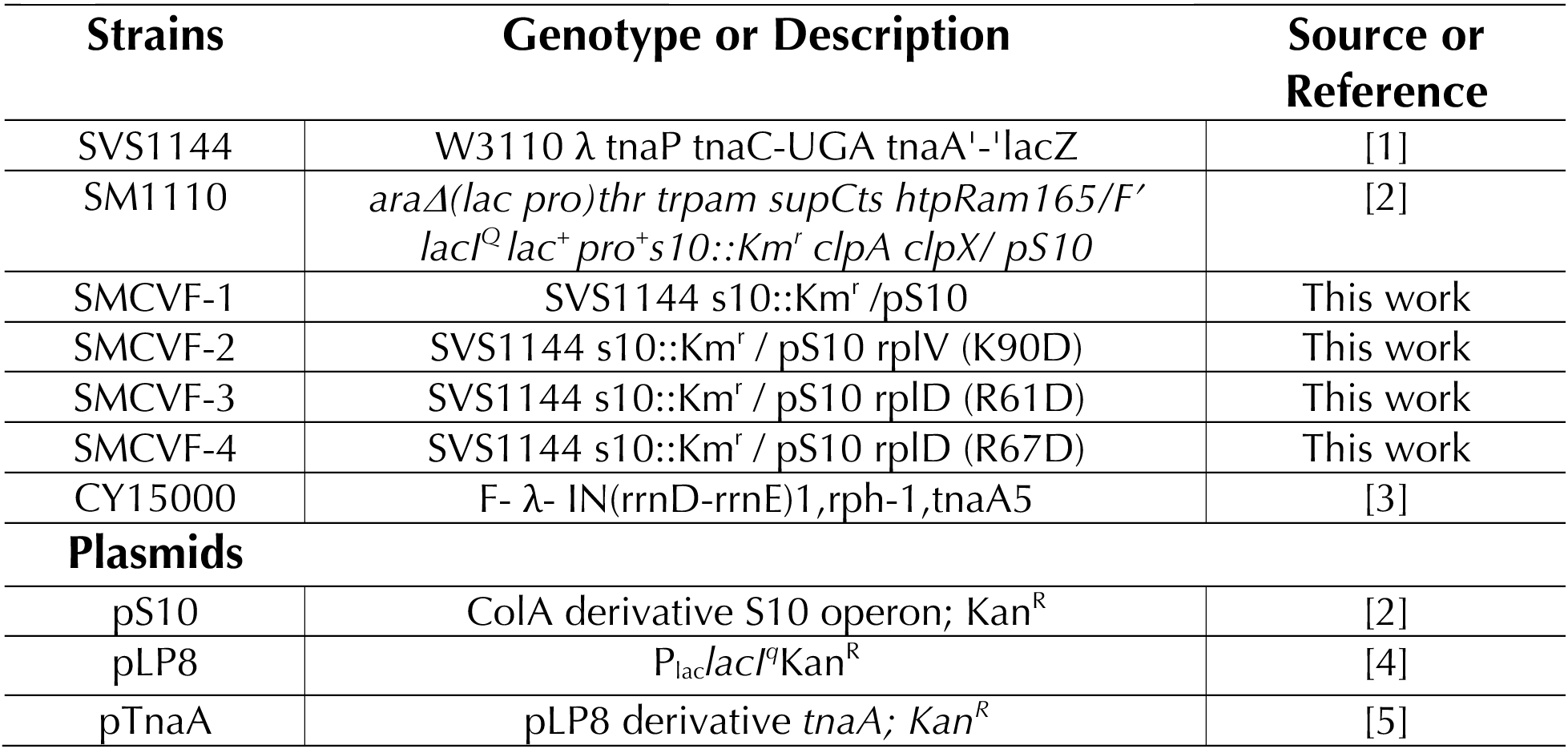
List of strains and plasmids used in this study

**Supplemental Table 2.**
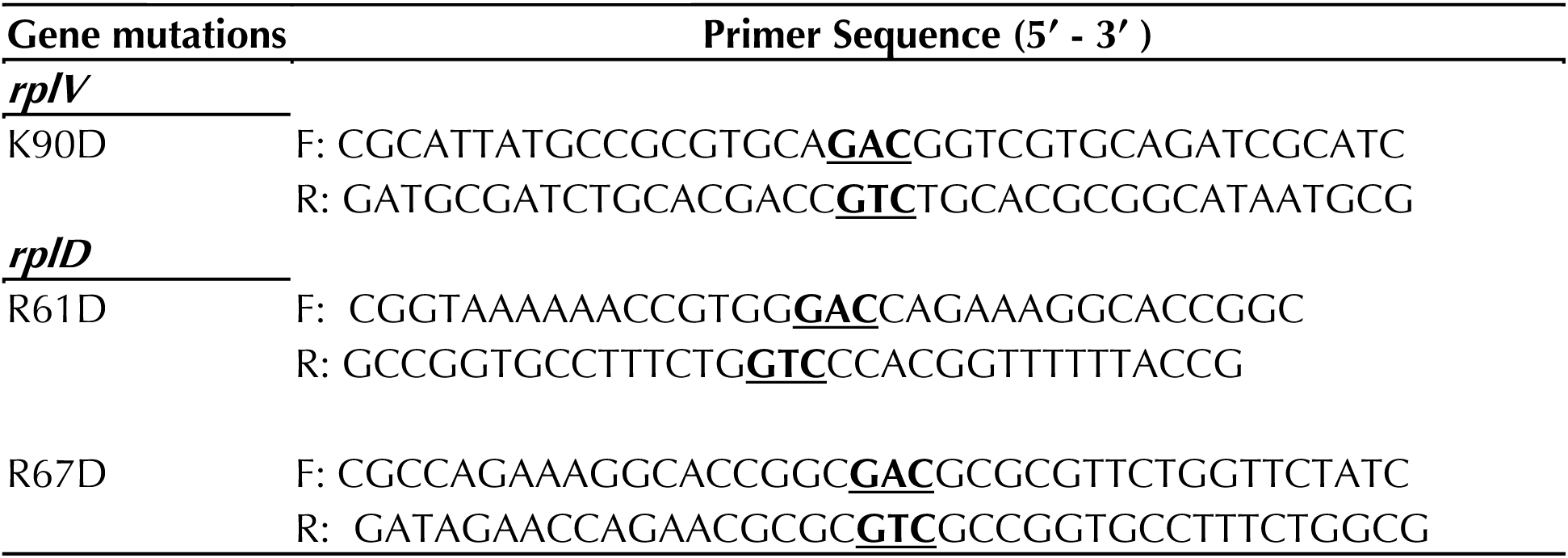
List of primers used in this study to mutagenize the proteins uL4 and uL22 produced from the pS10 plasmid

**Supplemental Figure 1.**
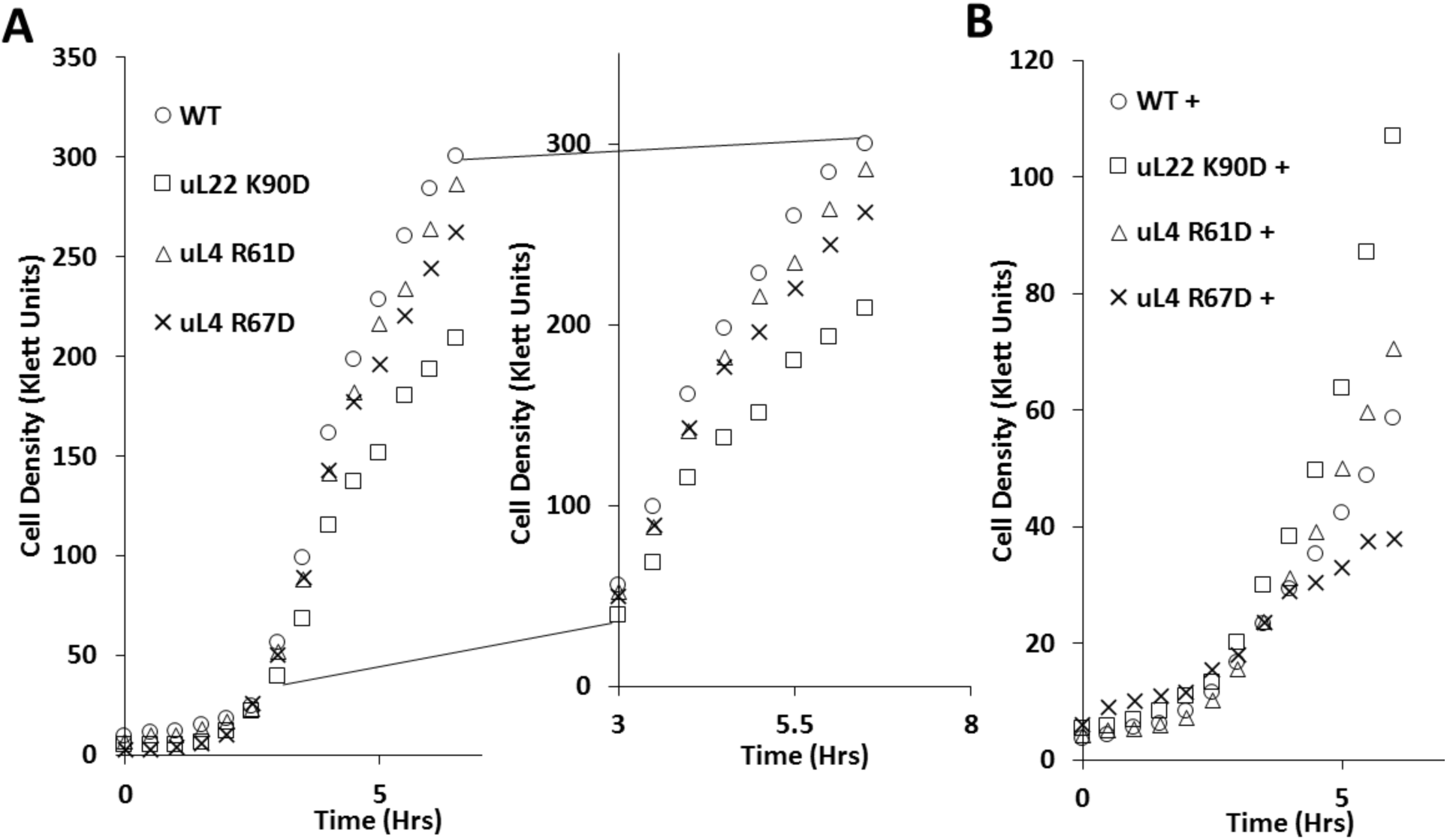
Cell growth of uL4 and uL22 wild type and mutant strains in rich media. Graphical representation of cell growth cultures monitored during the times indicated using a Klett nephelometer. (A) Right panel corresponds to a zoomed view of the resulting curve to highlight any growth differences in the late exponential and entry of the stationary phase. The uL4 mutants exhibit less differences in growth than the uL22 K90D mutant. (B) Cells were grown in LB supplemented with 73 µg/mL Erythromycin, indicated with a (+). All mutants outgrew the wild type except for the uL4 R67D strain. This figure is representative of at least two independent experiments.

**Supplemental Figure 2.**
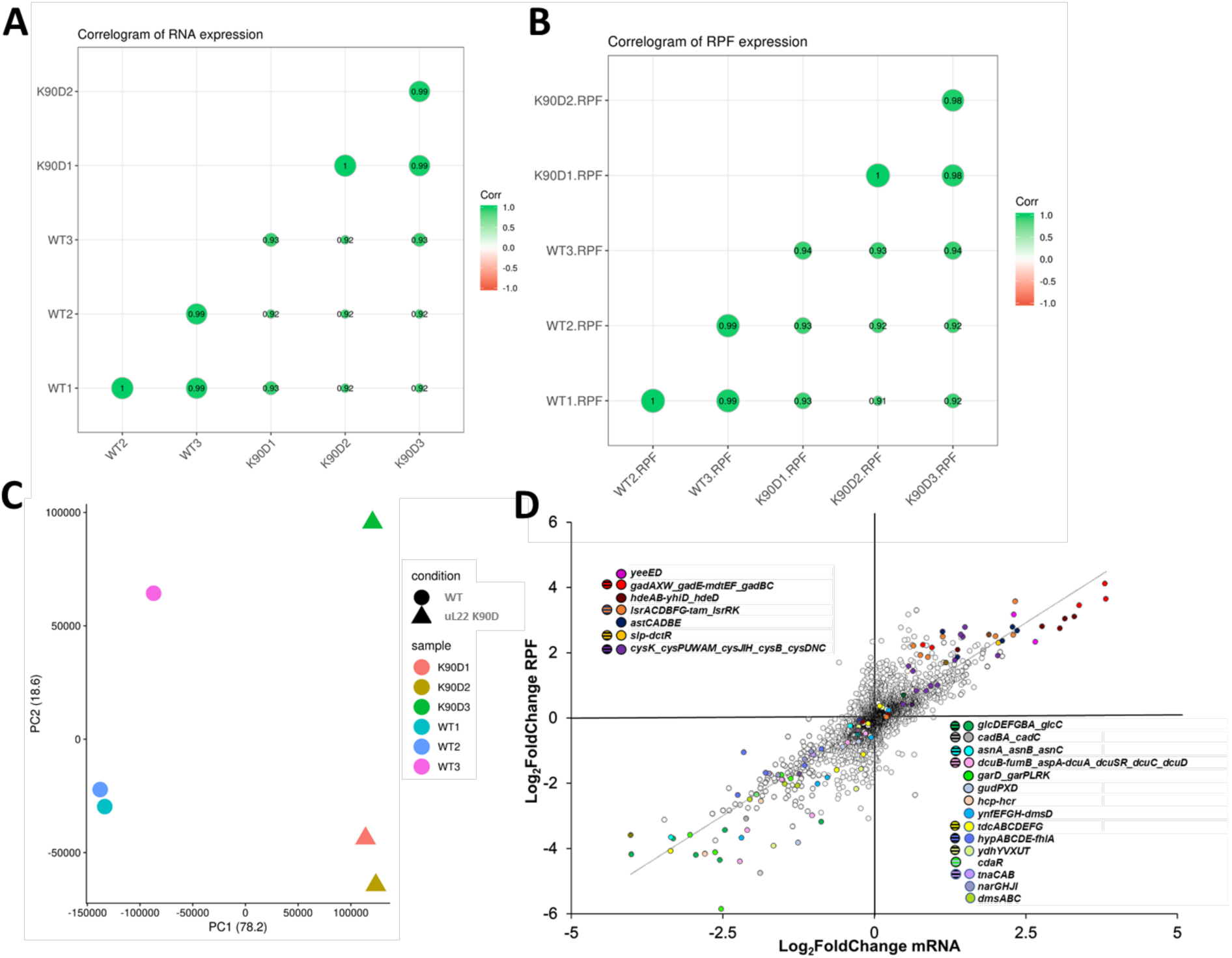
Reproducibility between ribosome profiling samples and correlation of RPFs to mRNA levels. (A) Correlograms of normalized RNA and (B) RPFs values between all samples. (C) Principal component (PC) analysis among different samples in two groups. For PC1 samples group according to the cell strain samples, and PC2 groupings correspond to different harvest dates of the samples. These groupings indicate true differences between the wild type and uL22 K90D cells. (D) Scatter plot showing changes in the values of ribosome protected fragments (RPF) per each gene are plotted vs their corresponding changes in mRNA levels. Genes corresponding to affected operons and their transcriptional regulatory factors are color coded. Operons shown at the top left quadrant showed high mRNA abundance and translation efficiency, whereas operons shown at the bottom right quadrant showed low mRNA abundance and translation efficiency in the K90D strain. Our data showed an R^2^ correlation of 0.66 between RPF and mRNA levels suggesting that the mRNA abundance was the major factor influencing translation of the majority of the tested genes.

**Supplemental Figure 3.**
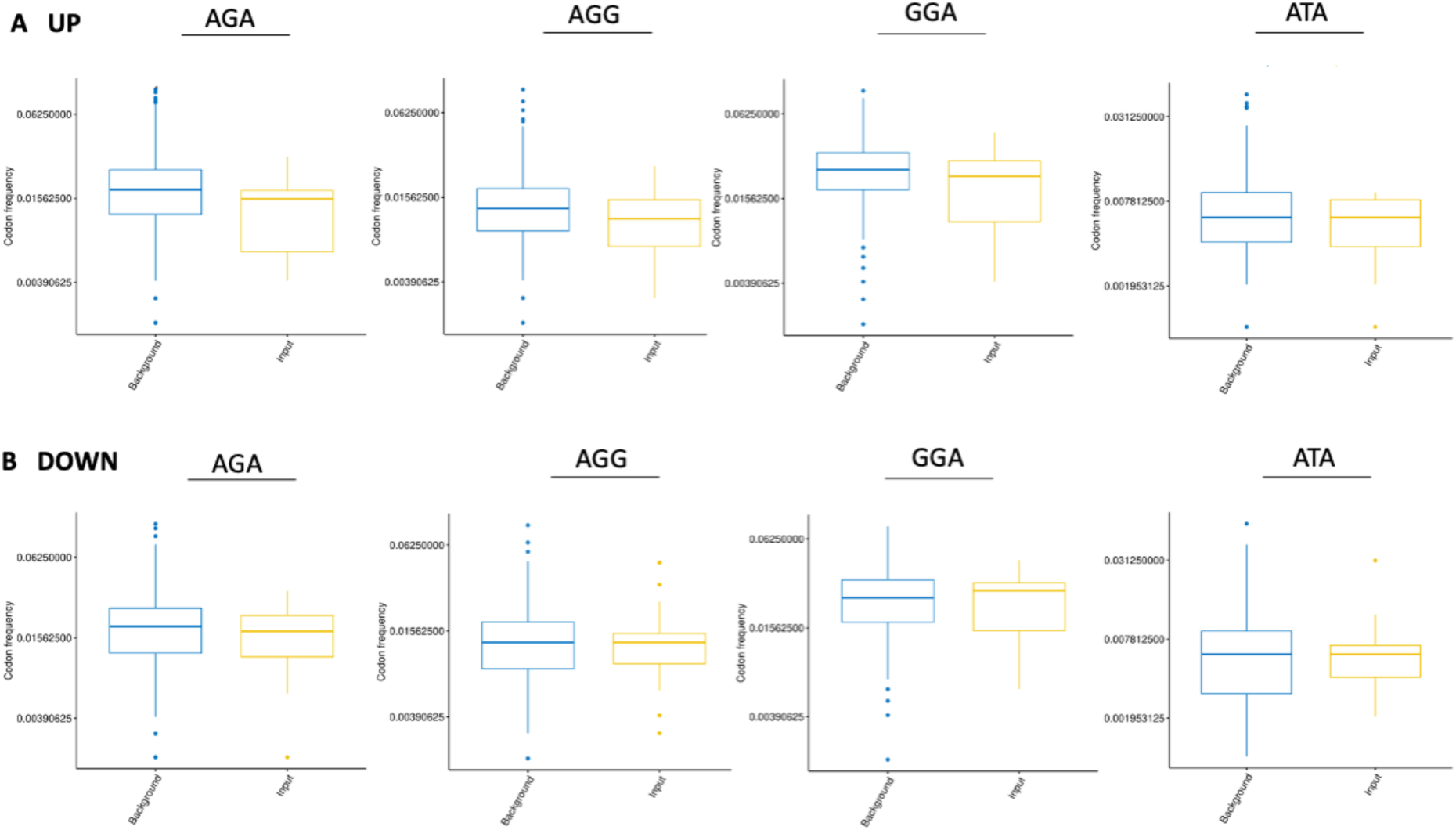
Comparison of rare codon frequency in the translationally up- or down-regulated genes in the K90D strain. Frequency of select codons in (A) genes showing high values of translational efficiency (up), and (B) genes showing low values of translational efficiency (down). Random genes are selected as background comparison.

**Supplemental Figure 4.**
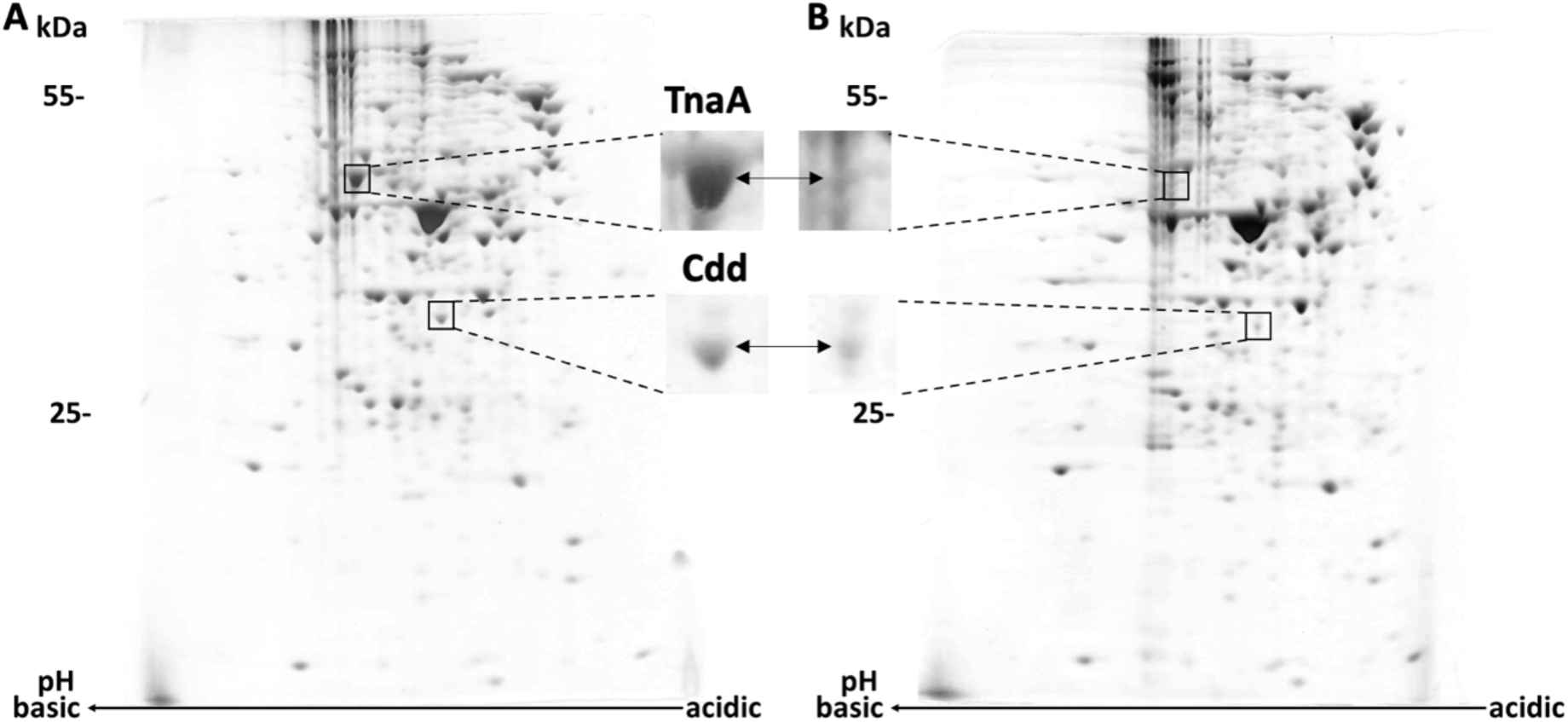
Analysis of the global expression of proteins of cells expressing the uL22 wild type and K90D mutant. Standard 2-D gel electrophoresis of cells containing (A) wild type (B) mutant protein uL22 K90D were performed as previously described [6]. Proteins were visualized using Coomassie Brilliant Blue R-250 stain. Zoomed comparison of tryptophanase (TnaA) and cytidine deaminase (Cdd), are shown. The intensity of the K90D spot was quantified as 59% for TnaA and 7% for Cdd that of the wild type. These values correspond with our ribo-seq data (see **Supplemental File**).

**Supplemental Figure 5.**
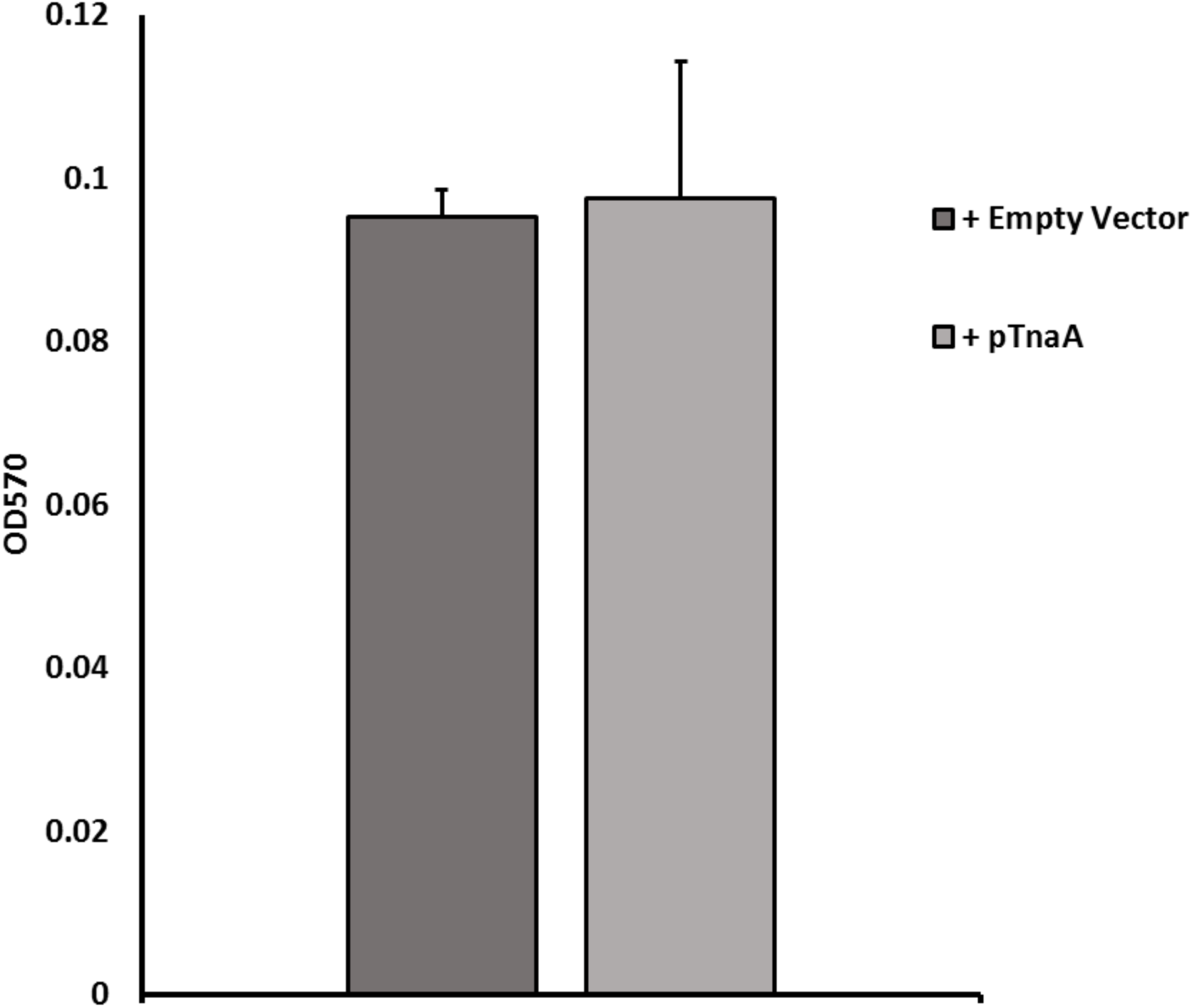
Biofilm formation in the presence or absence of tryptophanase in CY1500. An isogenic TnaA mutant (CY15000) was transformed with either pLP8 (empty vector) or pTnaA (pLP8 with *tnaA*) and biofilm formation of the resulting strains was measured. Results revealed that CY15000 biofilm formation is not affected by the overexpression of *tnaA*. Error bars were calculated using standard error between 6 biological replicates.

**Supplemental Figure 6.**
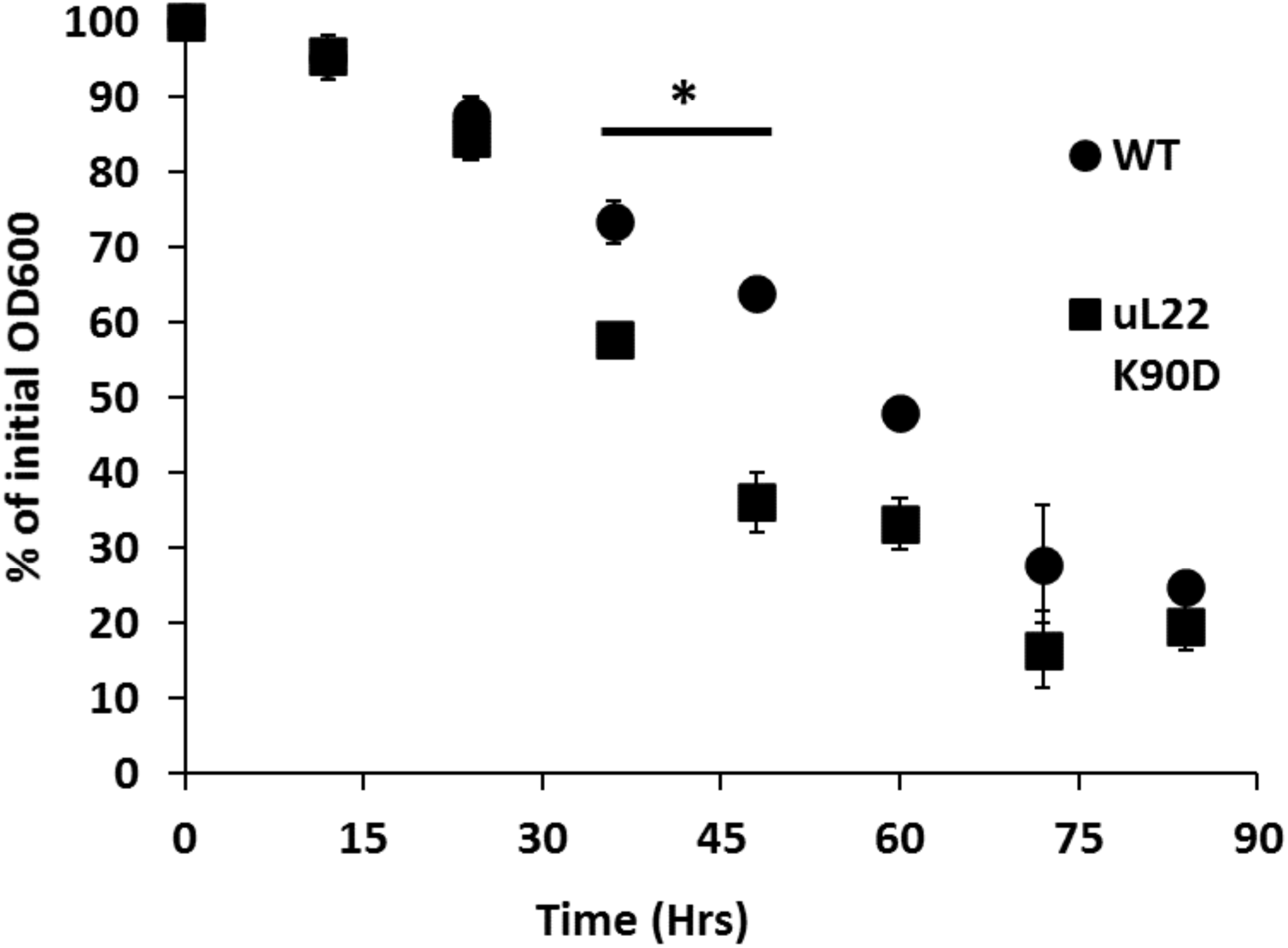
Sedimentation assay of cells containing wildtype or K90D mutant uL22 proteins. The sedimentation assays were performed as previously described [7] with the following modifications. Cells grew overnight in LB media at 37°C with shaking giving an initial OD_600_ of approximately 1. Cultures were then removed and placed on the bench-top at room temperature and in a static upright position for the duration of the experiment. The initial OD_600_ was recorded by taking a well-mixed sample from the upper liquid (inserting the pipet tip just below the surface of liquid) in the culture, and each subsequent sample was removed from the upper liquid of the culture every 12 hours for 84 hours. Data in the plot is expressed as percentage of initial OD. The mutant cells show significantly more sedimentation from time 36 (p-value= .03) to approximately 60 (p-value= .056) hrs. This data represents three independent experiments. Standard error is shown for each sample, and a student t-test was performed on the averages to determine statistical significance.

